# Kinetic patterns of single cell gene expression discriminate between the murine cellular responses to live attenuated and inactivated Yellow Fever vaccines

**DOI:** 10.1101/2024.11.26.625490

**Authors:** Budha Chatterjee, Christopher T. Boughter, Katrina Gorga, Yuko Ohta, Carly Blair, Elizabeth M. Hill, Zachary Fasana, Maxine Atuheirwe, Adedola Adebamowo, Farah Ammar, Natarajan Ayithan, JP Courneya, Ivan Kosik, Vel Murugan, Wilbur H. Chen, Marcela F. Pasetti, Martin Meier-Schellersheim, Nevil J Singh

## Abstract

The success of the live attenuated Yellow Fever vaccine (YF17D) that elicits immunity lasting over thirty years has made it a widely used model to understand the generation of durable protection. We compare the early single-cell level transcriptional response in mice to YF17D and an adjuvanted-inactivated, but less effective version (InYF). Within the first week, we identify 70 kinetic patterns in 45 cellular clusters, majority of which discriminate between the two formulations, some in a tissue and sex-specific manner. Intriguingly, differential transcripts fall into two categories, one whose association with YF17D or InYF is maintained even when decoupled from their cell-type of expression and the other where such cell-plus-gene pairing is critical to maintain differential marker status. We demonstrate applications of this resource, by identifying B cells with varied interferon and antigen responsiveness in relation to each vaccine. This high-resolution dataset is amenable to further biomarker discovery and hypothesis generation.

## Introduction

As new infectious agents emerge and mortality from previous ones continue to pose significant threats to human health, the need to rationally design new vaccines that can reliably engender life-long immunity remains significant^1-3^. Several clinical vaccines are currently administered to >70% of the world’s population and these reproducibly generate life-long immunity^4^. While these establish a goal, desirable in any new vaccine, recapitulating their efficacy has been challenging. For instance, even for multiple vaccines approved somewhat recently, correlates of protection wane within a few years after vaccination^5-8^. Unfortunately, vaccine development is still largely empirical. As a consequence, molecular mechanisms needed to engineer long-term efficacy remain unclear. One approach to resolve this issue involves reverse vaccinology, wherein successful vaccines are analyzed in mouse, non-human primate or human studies at a cellular and molecular level aiming to understand the precise pathways invoked *en route* to stable long term protective immunity. Immunizing model animals (such as mice) offers the opportunity to apply contemporary techniques that were not available when many of the highly successful vaccines were first licensed.

The Yellow Fever vaccine (strain YF17D), first developed in 1937^9^, with documented protection lasting 30+ years in humans^10,11^, has been particularly useful in this regard^12-17^. Insights gained from the fine-grained analysis of innate, B cell and T cell responses to this vaccine have already shaped our conceptual framework of immune response to effective vaccines^17^. Typically, these prior studies have extensively explored the effective, live attenuated YF17D, mostly using bulk-level techniques. Notably, viral-inactivation leads to a different version of the yellow fever vaccine^18-20^. Although inactivated vaccines can eliminate some rare adverse events, their typically poor durability often necessitates boosting, which can be demanding in economically constrained endemic areas^21^. In the context of reverse vaccinology, a high-resolution molecular comparison of an inactivated version with YF17D has the potential to map critical determinants of the host response that favor the development of durable immune protection.

Here, we leverage this well described vaccine-pair of live attenuated and inactivated YF17D to examine very early cellular responses in mice using single-cell RNA sequencing (scRNA-seq) in multiple immunologically relevant tissues. Rather than focusing on the known correlates of immunity (such as the B cell or T cell response at later timepoints), we sought to deeply probe early events that may discriminate live YF17D from adjuvanted inactivated YF (InYF) vaccination. Furthermore, in light of the emerging view that laboratory strains of specific pathogen free (SPF) mice have unprimed immune systems that affect subsequent vaccine responses^22,23^, our study first exposed mice to sequential infections with a virus (Influenza PR8) and a parasite (*Plasmodium yoelii* – followed by chloroquine cure) before administration of either YF17D or InYF. We also examined the responses in male and female mice separately to track the impact of sexual dimorphism^24-27^. The combination of single-cell level, tissue and sex-specific longitudinal analysis, along with the use of pre-exposed mice, offers a novel perspective on the underlying immunological differences triggered very early by these vaccines. Our approach yields data that can be a valuable resource to uncover previously unappreciated biomarkers discriminating between different formulations of vaccines, and, more generally, may eventually permit predicting vaccine efficacy at an early timepoint.

## Results

### Study design and data statistics

To comprehensively characterize longitudinal gene expression programs in response to the live-attenuated YF17D and inactivated YF adjuvanted with AS03 (InYF), we used single-cell preps from four immunologically relevant tissue sites (draining Lymph Nodes (dLN), irrelevant distal (non-draining) Lymph Nodes (ndLN), Spleen and peripheral blood (PBMC)) for CITE-seq analysis (Figure 1A, S1A-C, Methods). Previous reports suggest that SPF mice are poorer models of human immune responses compared to sequentially infected “dirty mice”^22,23,28,29^. Therefore, before vaccinations, mice underwent a *<u>V</u>*irus *I*mmune *<u>P</u>*lasmodium *<u>Ex</u>*posed (VIPEX) protocol (see methods) with sequential viral and parasite infections. This additional variable required using unchallenged VIPEX mice as controls, in addition to SPF (“naïve”) mice.

**Figure 1:**
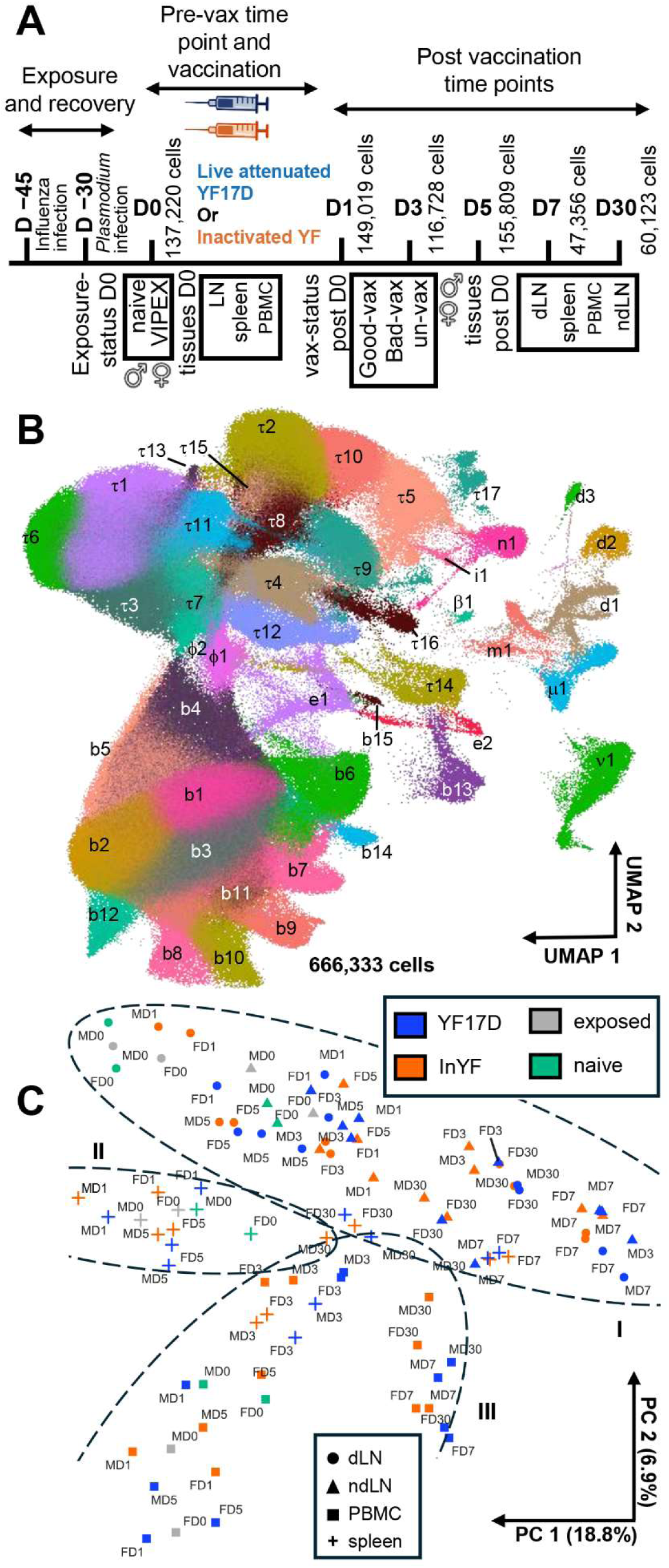
Clustering and principal component analysis of integrated data from all timepoints. (A) An overview of the experimental timeline. Mice were immunized on Day 0 (‘D0’) after receiving a flu infection 45 days previously (D −45) followed by a Plasmodium challenge 15 days later (D −30). Cell numbers analyzed on D0 and post-D0 timepoints, as well as, sex, vaccination status and tissues collected are listed below the scale. (B) UMAP of the integrated transcriptomic data from all cells analyzed in this report, divided into 45 major clusters. Each cluster is denoted with a symbol constituting of a letter followed by a number. τ → T cells, b → B cells, m → macrophage, μ → monocyte, n → NK cells, ν → neutrophil, β → basophil, d → dendritic cells, i → ILC, e → erythroid-like, ϕ → bi-phenotypic lymphocyte. (C) Principal component analysis of all samples, each averaged for the cell population within the sample (equivalent to bulk data). The ellipses I, II and III are manually sketched to indicate regions with samples from Lymph nodes, spleen and PBMC respectively. Each datapoint is labelled with sex (M or F) and time (indicated by D#) information.

Quality filtering, demultiplexing, normalization, integration, cell clustering and removal of singletons (from the full dataset submitted to NCBI) resulted in data from a total of 666,333 cells, analyzed here^30,31^ (Figure 1B, S1D, Methods). The clustered UMAP (Figure 1B, S2A) can be broadly separated into three overall regions, representing T cells, B cells and non-lymphocytes. This was facilitated by the antibody tags marking CD4, CD3, TCR β-chain, CD5 and CD8 for the T cell region, CD19 and MHC-II for B cells, and CD11b, Ly6G and CD11c associated with different classes of myeloid cells. To avoid arbitrarily assigning additional lineage subsets beyond these broad cell-types (to avoid potential confirmation bias with potentially similar canonical lineages) we assigned alpha-numeric symbols each cluster for all analysis (Figure 1B). The first character indicates cell-type identity: b being B-cells, τ (tau) T-cells, μ (mu) monocytes, m macrophages, ν (nu) neutrophils, n NK cells, d dendritic cells etc. The second, numeric character (e.g. d1-3) denotes sub-clusters (Tables S1A, B) and helps demarcate 45 distinct clusters in this report (Figure 1B). Although we do not assign specific sub-lineage labels the cluster markers included can be used for this, if necessary. For instance, b2, b4, b8, b11, b12 conforms to the follicular B cells and τ2, τ8, τ10, τ15 conforms to CD8s etc.(ImmGen annotation project^32,33^).

The 45 clusters captured the distributions of cells isolated from all four tissues examined in this study (Figures S2B-E, Tables S2A-D); but the number of cells present in each cluster varied with specific tissues. For instance, Spleen and PBMC contributed to a large proportion of the erythroid-like cells (e1, e2, 8944 of the 8987 cells in this cluster came from Spleen and PBMC), while the B cells in cluster b6 predominantly came from the spleen (19629 of 22099 cells). Similar tissue-specific differences were also found in several other clusters e.g. d2 (1804 of the 2399 total cells are of lymph node origin), τ12 (only 1053 of 14516 cells were from PBMC), τ9 (15453 of 16182 cells are from PBMC), b12 (almost exclusively from dLN with 5137 of 7476 cells).

### High-level analysis identifies dominant tissue specific expression profiles

Given these observations, we further explored cell densities across additional experimental covariates. Sex differences influenced frequencies of ν1 (neutrophils) with male mice having 2.2-fold higher numbers than females (arrowhead, Figure S2F). The “dirty-mice” protocol also had an impact; e.g. VIPEX mice harbored ∼3.4 fold higher number of cells in τ9, than naïve SPF mice (Figure S2G). Vaccination with InYF led to a more ν1 (3.3-fold) cells than YF17D, whereas YF17D elicited ∼4.1-fold more cells in b12 (Figure S2H). This extensive data allowed us to perform a high-resolution examination of multiple cellular dynamics in the early response with several differences in other clusters (e.g. clusters ν1, b12 and τ4) through time when projected across sex, tissue and vaccination status (arrowheads, Figure S2I).

As expected, principal component analysis (PCA) visualization of relationships between all the samples (Figure 1C) showed closeness of expression profiles in cells from same tissue. Tissue-specific properties tended separate less with the first two PCs even when from different timepoints or even vaccine formulations. A strong indicator of such proclivity is that samples from lymph nodes, irrespective of draining (dLN) or irrelevant (ndLN) node, all aggregate in the top half (oval region labelled ‘I’). PBMC are in the region labelled ‘III’ and spleen in ‘II’. The next best predictor of projected proximity was the timepoints. PBMC Samples at 1 day (D1), 3 days (D3) and 5 days (D5) after immunization were distributed along the “7 ‘o’ clock position (Fig 1C) compared to those from 7 days (D7) and 30 days after (D30). These relative distances were preserved in similar PCA using other layers of the data - surface markers, normalized RNA, or, all T-cells, all B-cells and all non-lymphocytes (Figure S3A-C).

### Contextualizing YF vaccine-pair single-cell data resource with the existing reports on live YF17D vaccine

Given previous studies analyzing responses to the YF17D vaccine using different approaches, we examined if the resource generated here encompasses and helps compare insights from those findings as well^12-17^. Querec et al (2006), focused on the impact of YF17D on antigen-presenting cells using bulk-RNA analyses and flow cytometry, to report markers such as CD80 and CD40 in CD11c^+^ dendritic cells (in dLN, spleen), IFN-γ in T cells (dLN) and TNF from DCs (PBMC)^12^. Indeed, we were able to map these primarily to DC cluster d1 and T cell cluster τ5 allowing for a granular dissection of the modified subsets (Figure 2A). Intriguingly, while previous studies compared SPF mice to those vaccinated with YF17D, the use of VIPEX mice also points out which changes are specific to YF17D over a bystander effect of a “first infection”.

**Figure 2:**
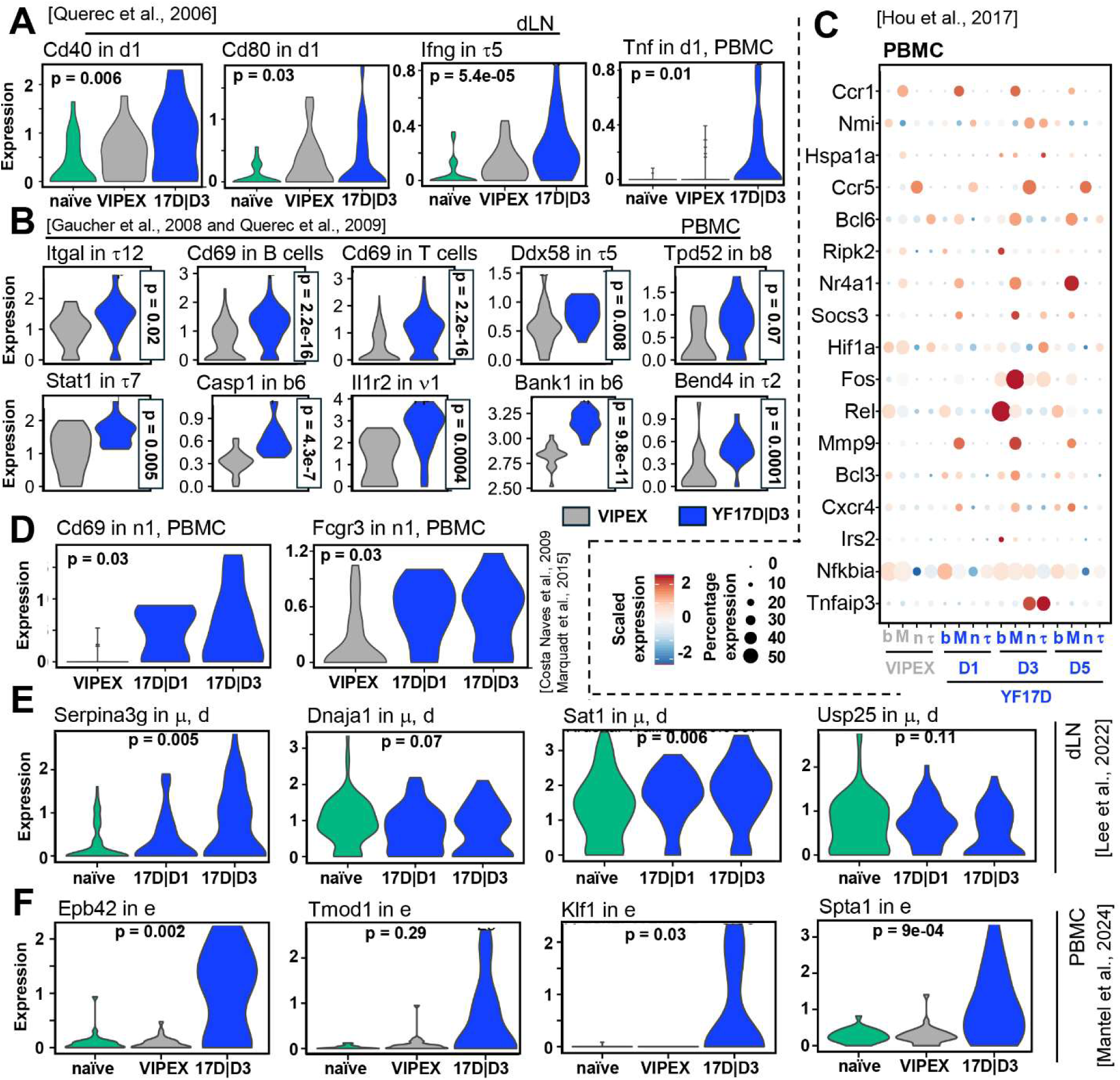
Assignment of cluster contexts for markers previously identified in the response to YF17D. (A) Expression levels, in our data, of CD40, CD80, IFNγ and TNF - markers originally reported in the Querec et al., 2006^12^ study. Values are extracted from the d1, d1, τ5 and d1 clusters. Tissue source is indicated at the top of the panels. YF17D is abbreviated as ‘17D’. Timepoint, D3, is marked, separated from 17D by vertical bar ‘|’. (B) Expression levels in mouse PBMC, of markers Itgal, Cd69, Ddx58, Tpd52, Stat1, Casp1, Il1r2, Bank1 and Bend4 (reported by Querec et al., 2009^34^ and Gaucher et al., 2008^35^ from human PBMCs). Specific cell clusters are indicated above each panel. (C) Dynamics of key genes reported by Hou et al 2017^36^ in our data on D1, D3 and D5, derived from a ‘pseudo-bulk’ analysis of all B cells (b), T cells (τ), Myeloid cells (M) and NK cells (n), from PBMC. (D) NK cell activation markers (Cd69 and Fcgr3 (CD16 in humans)) reported by Costa Naves et al., 2009^37^ and Marquadt et al., 2015^38^ evaluated in the n1-cluster of PBMC in response to YF17D. (E) Key markers from Lee et al., 2022^39^, Serpina3g, Dnaja1, Sat1, Usp25 extracted from naïve, D1 and D3, in monocytes and dendritic cells in this resource (F) Level of expression of genes (Epb42, Tmod1, Klf1 and Spta1) reported by Mantel et al., 2024^40^, plotted from erythroid-like clusters here. Name of the gene and cell cluster are the titles in each subplot. Two variable and three-variable comparisons were executed by Wilcoxon rank-sun tests and Kruskal-Wallis tests, respectively. P-values are mentioned on each plot as exact numbers.

Querec et al (2009) and Gaucher et al^34,35^ report CD69 upregulation in T and B cells, as well as anti-viral pathways (Stat1, Itgal, Ddx58) in PBMCs from YF17D vaccinated humans, using microarrays^34^. We also found Stat1 upregulation in several cell clusters. Some PBMC markers (BEND4, BANK1, TPD52) the authors predicted using machine learning algorithms, could also be validated in this dataset. The latter contemporaneous cohort study^35^ reported activation of inflammasome pathway by YF17D with induction of CASP1 and IL1R2. We observed the latter genes to be induced in b6 and ν1, respectively. All of these genes were induced on D3, compared to D0 VIPEX scenario, in PBMC (Figure 2B). Similarly, Hou et al^36^ investigated the kinetics of gene expression in a clinical setting over four weeks post-vaccination. To facilitate comparison with the observations in this bulk-level study, we calculated pseudo-bulk expressions from all B cells, all T cells, all myeloid cells and NK cells in current single-cell data. Most of the pathway hubs reported in this study could be validated from our sc-dataset (Figure 2C).

The early NK response is a characteristic of many anti-viral responses including YF17D ^37,38^. Indeed, CD69 and Fcgr3 upregulation in the n1 cell cluster of PBMC from D1 and D3 is also observed in our data (Figure 2D).

A recent study by Lee at al, which investigated immune response in draining lymph nodes in mice post YF17D vaccination at the single-cell level, focusing on myeloid cells, expectedly found a strong interferon signature again^39^. Apart from those, the authors also observed genes related to cell survival and viral infection, especially in monocytes and DCs. There were still important similarities in the data (Serpina3g, Sat1), and some differences (Dnaja1, Usp25). Expectedly, in our single-cell data, where VIPEX (not naïve) mice received YF17D, response of some of these genes were much less pronounced^23,28^ (Figure 2E). This highlights the importance of the previous exposure history; but the naïve cell signatures are available in this resource for comparisons.

Mantel et al, investigated temporal gene expression changes in PBMC of macaques after vaccination with live YF17D using bulk RNA-seq^40^. Apart from the expected antiviral pathway genes, and TLR-dependent dendritic cell activation, they reported pronounced activation of genes involved in heme-synthesis. We observed that these genes were induced in our erythroid-like (e) clusters on D3. Since the authors did not clarify whether the macaques were specific pathogen free (SPF) or not, we used data from both naïve and VIPEX mice for contextualizing their observation through our single cell data.

### Detailed cluster analysis identifies differences in the early cellular and transcriptomic response dynamics to YF17D vs InYF

Conventional flow cytometry-based analysis of immune cells showed relatively similar overall cellular dynamics between YF17D and InYF-vaccinated samples (Figure S3D, E). However, the higher resolution of the UMAPs allowed us to identify significant differences a (Figure 3, Table S2). A few myeloid clusters (m1, μ1, ν1) showed a surge in cell numbers on D1, in response to InYF. This increase was higher in InYF than YF17D vaccinated mice and also more pronounced in females than males (Figure 3B, panel ‘m1, D1’). Several T-cell clusters (e.g. τ1, τ3, τ8) elicited statistically significant depletion in response to InYF on D5 and D7, compared to VIPEX, in both sexes. At this early timepoint, vaccine-specific fluctuations in adaptive immune cells clusters could be noted. For instance, b12 and τ13 expanded significantly in response to YF17D and more in male mice (Figure 3B, panels ‘b12, D3’ and ‘τ13, D3’). In contrast, other B and T cell clusters (b8, τ4, τ14) were amplified to InYF on D5, to a comparable extent, in both sexes (Figure 3A, B, panel ‘b8, D5’). The dynamics in PBMC and spleen also showed significant differences but in fewer clusters and timepoints (Figures 3C and S4A)

**Figure 3:**
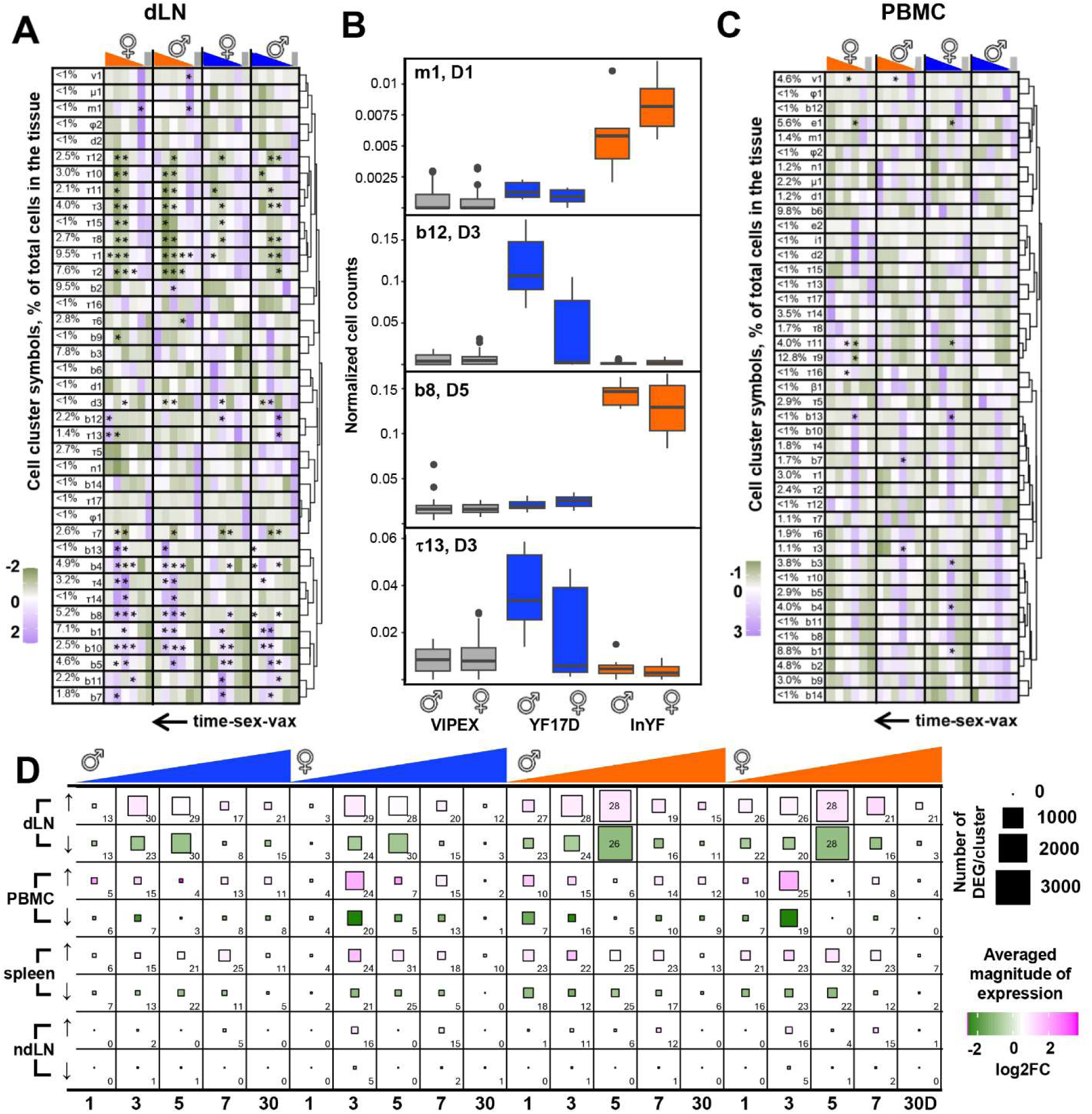
Global overview of cellular and transcriptomic dynamics to YF17D vs InYF within cell clusters. (A) The normalized number of cells recovered in each cell cluster, from each tissue and timepoint, are shown as a heatmap, arranged by hierarchical clustering. Columns (orange triangles = InYF and blue triangles = YF17D; ♀=Female, ♂=Male) incorporate color scaled lanes indicating density of cells on D1,3,5,7,30 (from right to left). Gray lanes at the end represent density of unvaccinated cells. The % values beside each cluster label (each row label, see column 1) indicates percentage of cells in that cluster across all time points, within all cells from that tissue. Clusters with less than 0.1% representation in a tissue are not shown. Asterisks represent statistical significance (adjusted p < 0.01) compared to the respective unvaccinated sex. Wilcoxon rank sum tests were executed on the cell numbers coming from different mice and different sequencing indexes, within the unvaccinated and vaccinated entries from a given cell cluster, followed by Bonferroni correction for multiple hypothesis testing. (B) Box plots of normalized cell numbers from representative clusters in dLN. See (A) for adjusted p-values. (C) Same as (A) in different tissue: PBMC. (D) Overview of differential gene expression (DEG) in this dataset, with each timepoint visualized down a column and tissues indicated by pairs of rows (top row in each pair = upregulated markers; bottom = downregulated). Samples are grouped by vaccine-type and sex as in A. Within the grid, the size of the squares (see scale key on right) represent the total number of DEG-cluster events (see text). Color scale filling the squares represents average overall expression for that group. Numbers below each column indicate total number of clusters where DEGs were detected. All comparisons are relative to exposed unvaccinated mice of the respective sex, pooled from multiple timepoints. DESeq2 algorithm^41^ was used for differential expression analysis along with Bonferroni FWER correction at adjusted p-value < 0.05, and log2FC > 0.25, or < -0.25.

To understand the extent of changes in gene expression on each timepoint post-D0, compared to unvaccinated (D0 and internal control VIPEX mice), we calculated differential gene expressions separately for male and female mice, who received either YF17D or InYF (Figure 3D, Methods). The distribution of aggregated counts for each cell cluster for a given timepoint, sex, and vaccine-type across sequencing samples were compared with those from the same cell cluster from unvaccinated, from the same sex, using the DESeq2 algorithm^41,42^.

Using sex and vaccine-type as discriminators at each timepoint across tissues (dLN, spleen, PBMC, ndLN), there were an average of 283 differentially expressed genes (DEG) across cell clusters (Figure 3D, median = 99 DEG-cluster events, read as ‘DEG from cluster’ events, see Methods). There were a large range for such DEG-cluster events: D5, dLN, ♂, InYF, down-regulated scenario had the largest number of DEG-cluster events: 3014, represented from 28 distinct cell clusters: 357 DEGs from b8, 122 DEGs from τ1 and so on. Whereas D5 and D30, PBMC, ♀, InYF, down-regulated scenarios had no DEG-cluster events. Differential expressions in non-draining lymph nodes were largely muted, as expected (bottom most pair of rows, Figure 3D). Overall, ∼58% of all DEGs was upregulated and the rest were downregulated, emphasizing that gene downregulation events can be important vaccination markers.

Longitudinally the largest number of DEG-cluster events were observed in dLN (69%), followed by PBMC (16%) and spleen (14%). The total number of DEG-cluster events is larger in response to InYF (67%), irrespective of sex, especially evident on D5 (Figure 3D, first two rows). Whereas overall DEG-cluster events across sex were only marginally higher in females (53%). Importantly, this large volume of DEG-cluster events across tissues, sexes and timepoints, detectable in our YF vaccine-pair single-cell dataset, suggests possibilities for mining novel ‘early’ markers of durable immune response to vaccination (Table S3).

Presenting the overlap of genes expressed in different tissue sites was plotted using Venn diagrams (Figure S4B) directly shows that DEGs across tissues were largely non-overlapping, as had been hinted at by the PCA. We next explored whether total number of DEG-cluster events were statistically associated with experimental covariates with Up-Set plots (Figure S4C, D). We observed that overall DEG-cluster sizes were associated with sex, vaccine and tissue, often with high statistical significance, across timepoints. Next, using Tree-Maps, we graphed trends in source cell clusters which represented most DEGs, over the covariates time, sex and vaccine (Figure S4E). These analyses showed that all covariates - tissue, sex, vaccine-type, timepoint and cluster identity had important influences on cell-type specific gene expression.

### Identification of transcriptomic biomarkers that distinguish between the vaccines as early as day 1 after vaccination

We analytically deconstructed the drivers of differential gene expression by examining the sources of maximal variation in the overall single-cell dataset using variance decomposition analysis (VDA)^43,44^ independently for each responding tissue (Figure 4A, B, S5B). Sex explained >50% of variation for a handful of well-known sex-linked genes while time explained >50% variation for 42 genes in PBMC, but only 10 genes in dLN. Importantly, without cluster information, the vaccination-status (unvaccinated, YF17D, InYF) was not a sufficient covariate to explain >50% of variation in any feature. Adding cell cluster information to vaccination-status was crucial and as a conjugate covariate, it explained >50% variation in 36 genes in PBMC and 15 genes dLN. Combining cluster, vaccine-type and sex covariates explained >50% variation in 64 genes in PBMC. Further, as we created cluster-vaccine-time conjugate covariate, that explained >50% variation in 108 genes in PBMC, 152 genes in spleen and 186 genes in dLN. This indicated that marker-discovery for vaccine-efficacy should incorporate covariates, including the source cell-type of DEGs. As reported in several other single-cell datasets^43,45^, a large part of the variation in gene expression could not be explained by the covariates (‘Residual’, Figure 4A, B).

**Figure 4:**
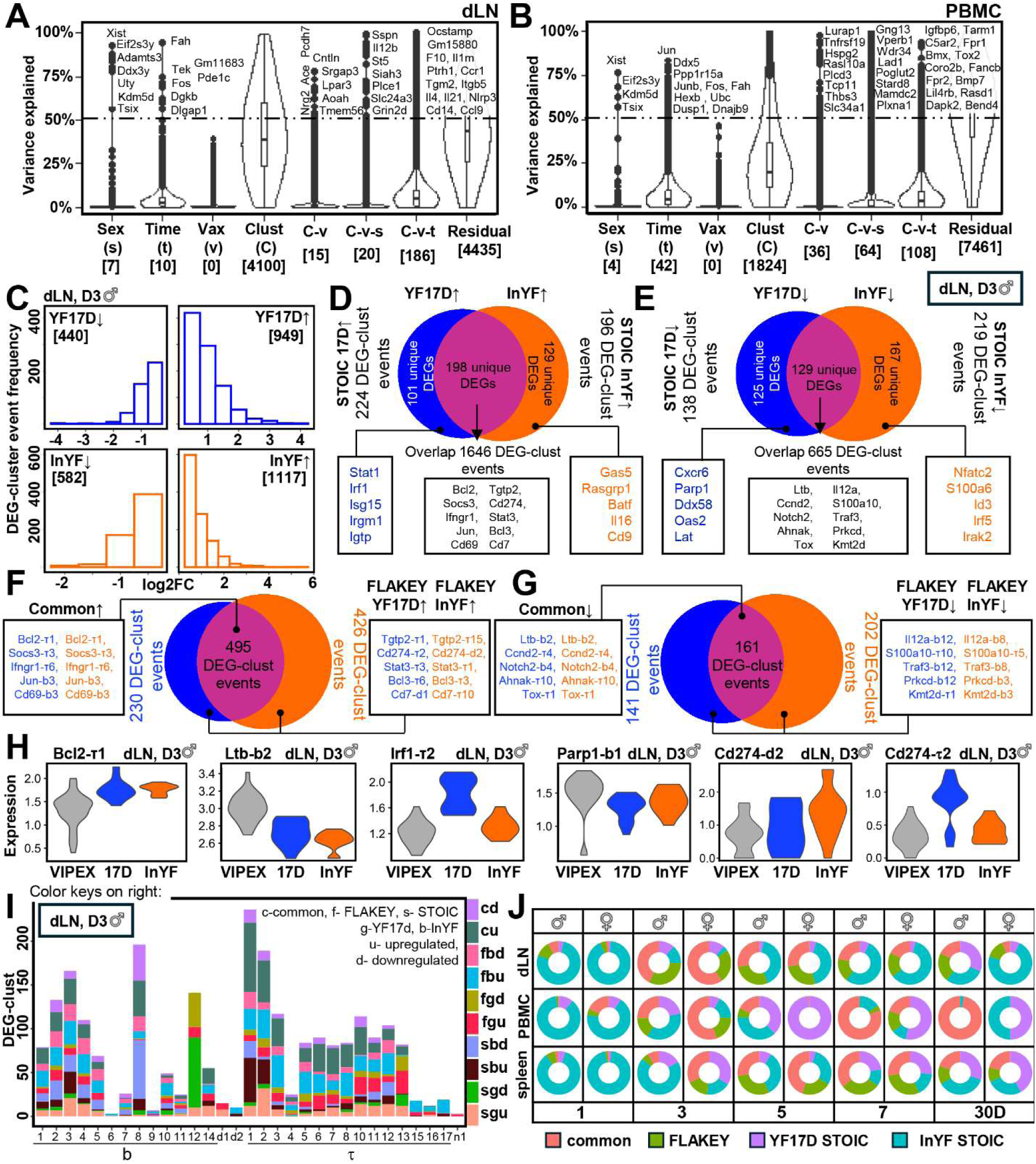
Categorization of markers associated with different experimental variables. (A, B) Representation of the contribution of each covariate or combinations thereof, towards the differential expression observed, by Variance decomposition analysis. dLN (A) and PBMC (B) for D0, 1, 3, 5. Covariates are listed on the X-axis. Numbers in square brackets are the number of genes for which the corresponding covariate explains more than 50% of the variation (dashed line). C-v: Cluster-ID + vax, C-v-s: Cluster-ID + vax + sex, C-v-t: Cluster-ID + vax + time. (C) Distributions of binary log fold changes in differentially expressed genes (DEGs). The D3 timepoint from ♂ dLN is chosen as a representative analysis (additional times/tissues are in Figure S5A, Table S4). (D, E) Venn diagrams of DEGs upregulated (D) or downregulated (E) relative to VIPEX mice. Data shown is of D3|♂|dLN data. The genes unique to each vaccine treatment are categorized as STOIC (see Text), compared to those shared by both responses (YF17D Ո InYF). Boxes below highlight a few of the exclusive and overlapping genes. (F, G) The genes found in the shared category were re-plotted, but with the relevant source cluster-of-expression information included. This identifies a new category of genes (FLAKEY, see text) which, when associated with source cluster-ID, is unique to YF17D (Blue) or InYF (Orange). The intersection identifies the Common signatures elicited by both. Names in boxes illustrate a few examples of Genes and their cluster (hyphenated). (H) Violin plots of selected DEG-cluster events listed in (C-G) for illustration with gene name & hyphenated cluster-ID indicated on the panel title. Gray: VIPEX, blue: YF17D, orange: InYF. (I) Distribution of STOIC, FLAKEY and common genes within cell clusters for D3 ♂ dLN (Table S4, Figure S5A for other tissues, timepoints, sex). (J) Donut plots depicting the distributions of STOIC, FLAKEY and common genes across all tissues, timepoints and sexes

Subsequently, we explored gene expression markers linked to YF17D or InYF in the responding tissues, i.e., dLN, spleen and PBMC, independently. Given the size of our dataset, we are using one slice of the data – i.e. from D3, ♂, dLN DEGs in the representative figure (Fig. 4C), with the distributions of upregulated (↑) or downregulated (↓) DEG-cluster events in each vaccine, relative to the VIPEX samples. Unlinking the names of the genes from their cluster-ID (e.g. Tgtp2 from τ1 and Tgtp2 from τ15 would be reduced to just Tgtp2), we found that there were specific genes that consistently associated with either YF17D or InYF (upregulated, Fig 4D and downregulated, Fig 4E, relative to VIPEX). The expression of the genes as markers for each vaccine therefore do not require association with a cell cluster and we refer to them as STOIC (<u>S</u>table <u>T</u>ranscriptomic <u>O</u>bservation Invariant to <u>C</u>lusters). A large fraction of the genes, falling in the overlap of the Venn circles in Figures 4D and 4E could further be subdivided into two categories by restoring their cluster identity. In some cases, restoration of cluster identities where the genes are expressed, showed that they were commonly upregulated or downregulated (relative to VIPEX) and these are labeled as “Common” events (Figure 4F, G, callout boxes originating from the intersections). Interestingly, the remaining shared genes, upon restoring their cluster-associations were found to be differential features between YF17D and InYF. We denoted these as FLAKEY genes (<u>FL</u>uctuating <u>A</u>ssociation to <u>KEY</u> treatment’, Figure 4F, G). These mutually exclusive gene groups, i.e., STOIC, FLAKEY and common together encompass all DEG-cluster events for the vaccine-pair, for a given timepoint, sex and tissue (Figure S5C).

The STOIC genes are robust marker candidates, since they are exclusive for either of the vaccines (Figure 4D, E, S5A, Table S4). A STOIC event can be exclusively YF17D ↑, YF17D ↓, InYF ↑ or InYF ↓. They cover ∼25% of all DEG-cluster events in D3, ♂, dLN data. For instance, Irf1 from τ2 was one among 224 STOIC YF17D ↑ events (Figure 4H, ‘Irf1’ panel). On the other hand, Parp1-b1 was one among 196 STOIC InYF ↑ events, for D3, ♂, dLN scenario (Figure 4H, ‘Parp1’ panel).

FLAKEY events could be easily missed in bulk analysis where the cluster-ID from single-cell evaluation is missing. In the D3, ♂, dLN data we found 230, 426, 141, and 202 FLAKEY YF17D ↑, YF17D ↓, InYF ↑ and InYF ↓ events respectively. They constituted ∼35% of DEG-cluster events in D3, ♂, dLN data. For an illustrative example, Cd274 (PDL1 in humans) gene is upregulated in response to InYF in cluster d2, but in response to YF17D, Cd274 is upregulated in some specific T cell clusters, τ2, τ7, τ10 (Fig 4H, ‘Cd274’ panels).

In D3, ♂, dLN there were 495 and 161 common upregulated and downregulated events, respectively (Figure 4B, C, S5A). Interestingly, more than 20% of all cluster specific DEGs were common between YF17D and InYF (∼16% upregulated, ∼5% downregulated). For instance, Bcl2-τ1 and Ltb-b2 are similarly upregulated and downregulated, respectively, in both vaccine types (Figure 4H, ‘Bcl2’ and ‘Ltb’ panels).

The contribution of each cluster to STOIC, FLAKEY and common genes in the D3, ♂, dLN (Figure 4I) varied, with τ1, having the highest number of common↑ events (79 events). Among B cells, b8 harbored the largest number of STOIC events downregulated in InYF (67 events). In contrast, b12 constituted an unusually large number of FLAKEY↓ and STOIC↓ events both in response to YF17D (39 and 80 events, respectively). Very low representations, as seen from b6, b9 and n1 are expected, since they are essentially absent in dLN (Figure 2B-E).

Next, we carried out this characterization of common, STOIC and FLAKEY genes across all timepoint-sex-vaccine scenarios (Figure 4J, Table S4). D1 was replete with STOIC events responding to the InYF. D3 and D5 timepoints had higher representation of common genes, as well as FLAKEY genes. Note that genes may change their identities over time, sex and tissue, i.e., over the remaining covariates. For example, a gene can be FLAKEY on D5 but STOIC on D1, in a given tissue, sex (e.g., Tnfaip3, dLN, ♂).

### Kinetic pattern search reveals subsets of genes in specific clusters that follow distinct patterns of expression over time in response to YF17D vs InYF

Given the multiple, close timepoints available to analyze the early phase of the response, we then explored kinetic patterns in the gene expression in cell clusters, while keeping all covariate information intact. Transcriptome-level exploration of kinetic patterns of gene expression can reveal shared drivers and tissue-specific variables such as inflammation, cytokine signaling etc. Significant over-representation of a kinetic pattern of gene expression in a given cell-cluster, compared to other cell-clusters, should be indicative of concerted, cell-type specific, immuno-biological processes. Any DEG-cluster event which appeared at least once in our DGE analysis (Figure 3D) was included in the kinetic pattern analysis.

Using partition around medoid (PAM) algorithm^46^ (Methods, Figures S5C, D), we first isolated kinetic patterns in the dLN layer. We detected 30 different patterns (Figure S5E, Table S5A), four of which are highlighted for illustration here (Figures 5A, S5E, star-marked, pattern id: LNVI, LNVII, LNXIII, LNXXI). Pattern LNVI was associated with τ6, τ13, d1, d2. Example transcripts, Fcer1g and Plec were upregulated in d1 on D1, in response to InYF. The signature of TLR activation was also evident in d1. On the other hand, several genes in LNVI:d2 (significant association between a kinetic pattern and a cell-cluster is presented as patternID:clusterID), such as Actr3, Wdr1, s100a10 were classically related to actin metabolism and different cell-repair mechanisms. LNVII was associated with b8, a delayed downregulation pattern with InYF, on D5. Classical NF-κB target genes such as Bcl10, Cd40, Tnfα were present in LNVII:b8.

**Figure 5:**
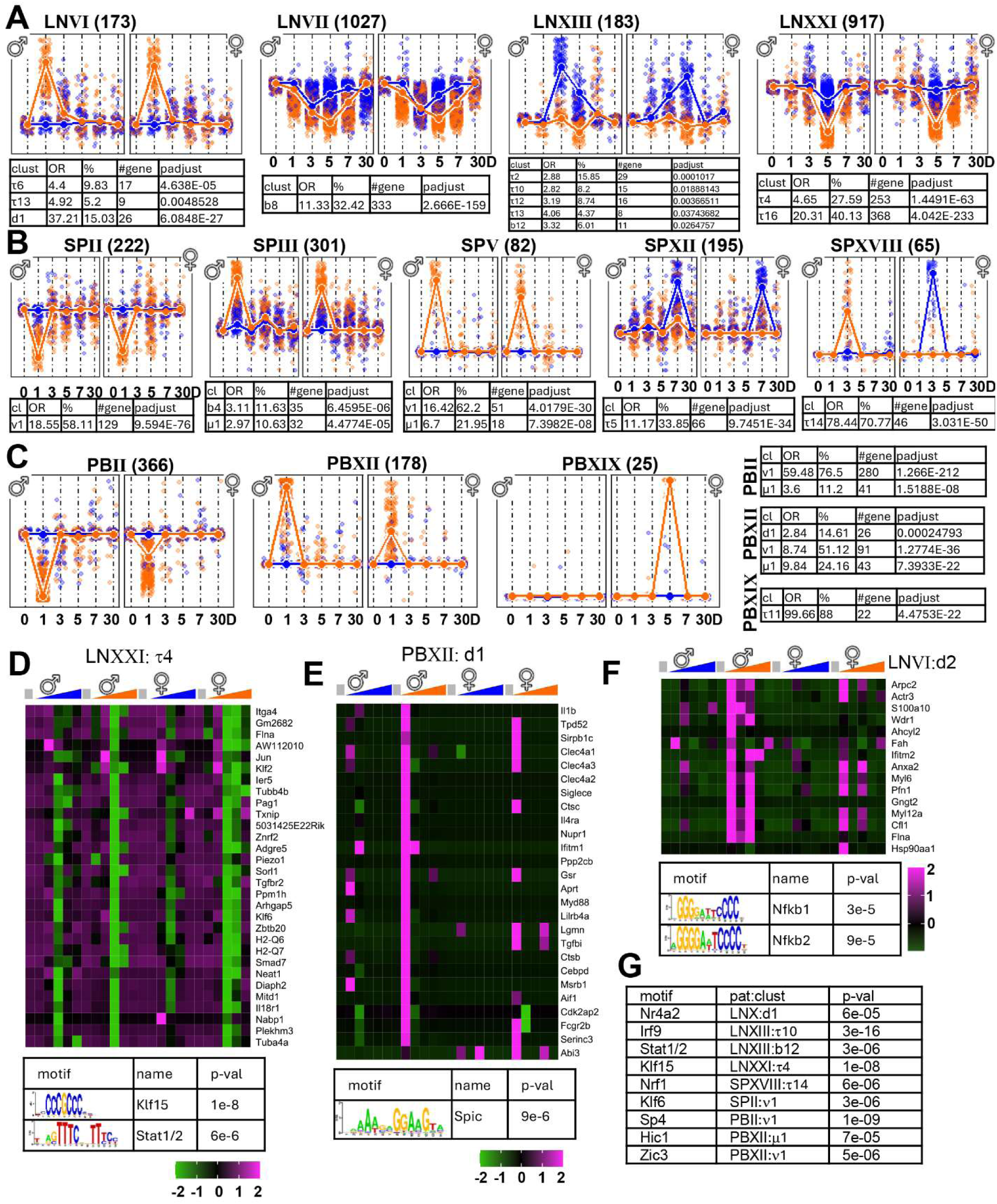
Kinetic patterns of gene expression from specific cell clusters distinguish early responses to YF17D vs InYF. (A) Representative outputs from kinetic clustering analysis showing 4 of the discriminating patterns found in dLN. Each panel shows patterns in ♂ (left) and ♀ (right) as sub-plots, over D0, 1, 3, 5, 7 and 30 (X-axis). Dots in the plot are standardized fold changes from individual DEG-cluster events within a given pattern and bold lines represent the medoid event among of these. InYF = Orange; YF1D = blue. Numbers next to the label (title) for each pattern represent the number of DEG-cluster events following that pattern. Boxes below the patterns are the results of one tailed Fisher’s exact tests, indicating positive associations between cell clusters and kinetic patterns. P-values were further adjusted with Bonferroni correction. OR: Odds ratio. (B) Similar to A, representative discriminating patterns from spleen and (C) PBMC. (D) Enrichment of transcriptional motifs in pattern-cluster associations. Heatmaps of gene expression and corresponding motifs below the heatmap. Scales below represent fold change, saturated at (-2, 2) for better visualization. DEG-cluster events from τ4, following the pattern id LNXXI. (E) DEG-cluster events from cluster d1 following pattern id PBXII. (F) DEG-cluster events from cluster d2, following pattern LNVI. (G) A brief list of motifs enriched in specific pattern:cluster associations (see Table S6 for a complete list).

LNXIII was closely associated with interferon-β response, specifically against the YF17D. Interestingly, the peak in LNXIII was on D3 for males but on D5 for females. Representation of this type-II interferon response from several T cell clusters and b12, indicates that lymphoid cells actively participate in such response. LNXXI again showed a pattern of downregulation on D5, for the InYF. The pattern was somewhat muted in females. In LNXXI:τ16 several circadian rhythm related genes, such as, Clock, Cry1, Ncor1 were overrepresented. An enrichment of genes participating in progesterone signaling pathway, such as, Tref1, Ubr5, Nedd4 was also observed. LNXXI:τ4, in contrast, showed downregulation of innate activation markers Cd86 and Ddx60.

Similarly, we found 26 distinct patterns in the splenic response (Figure 5B, S6A, Table S5B) of which five are illustrated (Figure 5B, ‘star-marked’ in S6A). The ν1 and μ1 clusters contributed to many of these and correspond to antiviral responses (e.g. Ifitm1, Apobec3 etc.) but others such as SPXII:τ5 highlight parallel changes in T cell migration and adhesion (e.g. Itga4, Ccl5, Itgb1, Coro1). Pattern SPXVIII was distinct in the response to InYF in males but in females it switched prominence to YF17D. Several of these genes in SPXVIII:τ14 were functionally related to cell division process: Prc1, Ccnb2, Fbxo1 etc.

In the PBMC layer we describe 20 patterns – where many had sex-specific trends (Figure 5C, S6B, Table S5C). Three patterns PBII, PBXII and PBXIX, were overrepresented by DEG-cluster events from ν1 and μ1, as was the case with spleen. In PBII:ν1, Tgfb1 and Cd4, Cd8 were downregulated on D1 in response to InYF. Similarly, In PBII:μ1, chemokine production (Cx3cr1) and overall innate response were muted in response to InYF. In PBXII:d1 several MHC genes were upregulated on D1, and they produced Il1b and Il4ra, specifically in response to InYF. PBXIX:τ11 on the other hand, expressed genes related to platelet degranulation: Lyn, Syk, as well as markers of differentiation Cd74, specifically in females, in response to InYF.

Since kinetic patterns may indicate concerted temporal gene expression, we examined the enrichment of transcription factor motifs^47^ in the promoters of the genes within pattern:cluster associations. There were statistically significant associations of known transcription factor motifs in almost every instance (Figures 5D-G, Table S6). For instance, the LNXXI pattern of delayed repression in τ4, was associated with Klf15 and Stat motifs (Figure 5D), whereas the early upregulation in LNVI pattern in d2 was associated with NF-κB motifs (Figure 5F). The complete charts of pattern:cluster associations and transcription factor enrichments are available as supplementary data (Figure S7A-C, Table S6).

### Bayesian networks reveal distinct communities of DEG-cluster events for YF17D and InYF

Gaining mechanistic insights from large-scale biological data is an ongoing challenge^48,49^. To illustrate that the current data resource is amenable to such analysis, we constructed Bayesian networks using the sex-agnostic DEG-cluster events for a given timepoint and vaccine, from the dLN layer. The granularity of our data, even after ‘pseudo-bulk’ing (separate cohorts for males and females, multiple replicates for each animal, see Methods), enabled us to successfully carry out these calculations (Figure 6A-C, S7D, E).

**Figure 6:**
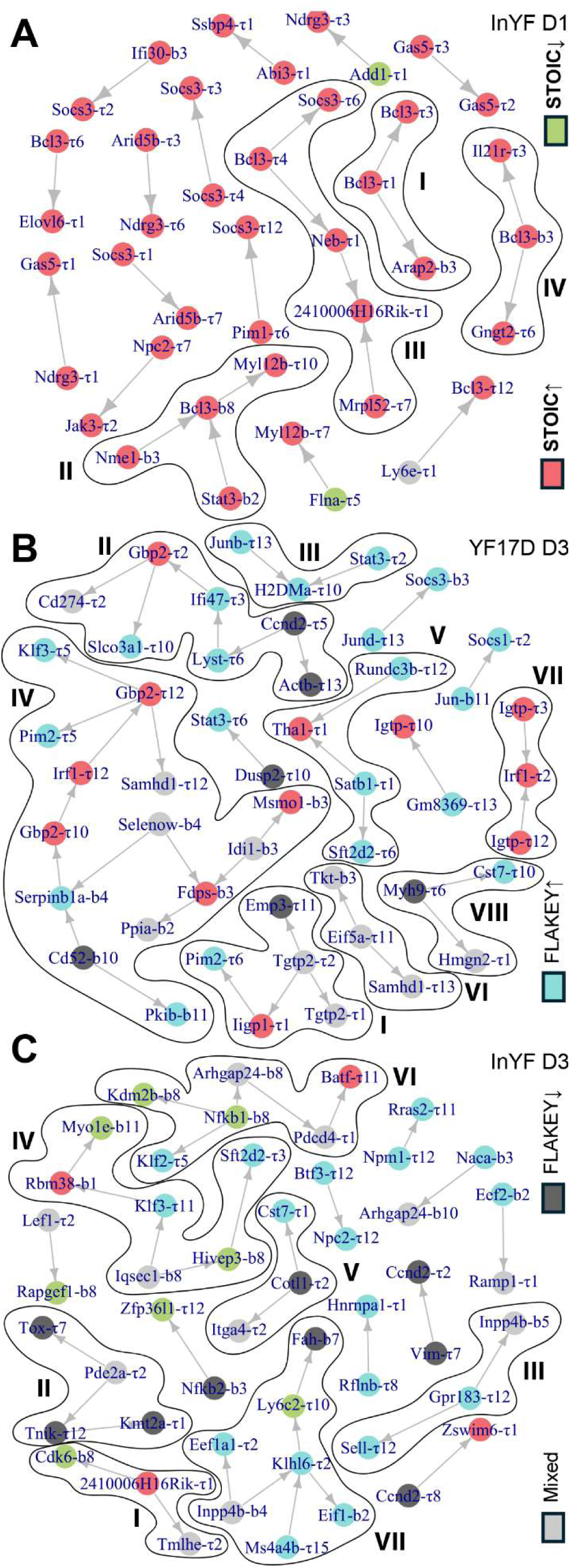
Bayesian structure learning of relationship characteristics discriminating responses to YF17D and InYF. (A) DEG-cluster events found in both males and females on D1 in response to InYF were used as nodes to construct conditional independence networks (see text, Methods). The nodes are color coded according to their gene group identity. When such identity is not identical in males and females, it is designated as ‘mixed’. Communities with three or more nodes are marked with Roman numerals. (B) Same as (A); on D3 in response to YF17D. (C) on D3 in response to InYF.

In these networks, an edge indicates a correlation between the two nodes (pseudo-bulk samples for a given DEG-cluster event), which is independent of other nodes in the network. The direction of the edges are suggestive of causation, iteratively assigned by Grow-Shrink Bayesian structure learning algorithm^50^.

Response to YF17D on D1 was relatively muted in dLN, compared to that to InYF. We could construct a network only for InYF, and not YF17D, on D1 (Figure 6A). Activation of classical NF-κB target genes such as, Bcl3 was observed in several T cell and B cell clusters, which potentially influenced the expression of other genes such as Socs3, Arap2, Il21r.

Networks on D3 could be constructed for both YF17D and InYF (Figure 6B, C). The largest community in YF17D D3 network (community IV) was enriched with several genes from metabolic pathways of cholesterol biosynthesis (Msmo1, Idi1, Fdps) and oxidoreductive stress management (Ppia, Selenow) from B-cell clusters and antiviral (Gbp2, Irf1, Samdh1) response from T-cell clusters. While the post-immunization involvement of sterol metabolism in B cells^51,52^, and oxidative stress post-YF17D vaccination^53^ are not unknown, yet their potentially close crosstalk with T cell subsets is under-appreciated. Among other novel communities, the nexus between Cd273 (PDL1), antiviral response genes (Gbp2, Ifi47) and cell cycle regulators (Ccnd2) across T cell subsets was provocative (Figure 6B, community II).

The signaling milieu of InYF D3 (Figure 6C) was strikingly disparate from YF17D D3. The communities appeared to be driven by NF-κB (as observed in D1 network, Figure 6A), Ras and phosphoinositide signaling. D5 networks of either vaccine indicated a plethora of less known, potentially causal connections (Figure S7D, E).

### Integration of biomarker discovery with kinetic patterns in the cell clusters permits physiological interpretation of early immune responses

The resolution and density of the YF vaccine-pair sc-dataset analyzed here, offers extensive resources for biological interpretation. To illustrate this utility, we focused on the differential dynamics within b8 and b12, two of the B cell clusters that showed significant density differences between YF and InYF (see Figure 3). Although we avoid computationally assigning specific sub-lineages here, key molecules represented here can be used to infer the developmental status of many cells herein. For instance, the reduced expression of Pax5 in b8 relative to b12 (Figure 7B), with increased expression of Xbp1 in a segment of b8, indicates that this includes activated B cell populations further committed to a plasma cell fate than b12^54,55^ but neither are fully differentiated plasma cells (as indicated by expression of Fcer2a and Ms4a1, which per the ImmGen database^32^ are heavily downregulated in FACS-sorted PCs). The upregulation of MHC and CD86 in b12 is consistent with a later follicular or early GC stage cell.

**Figure 7:**
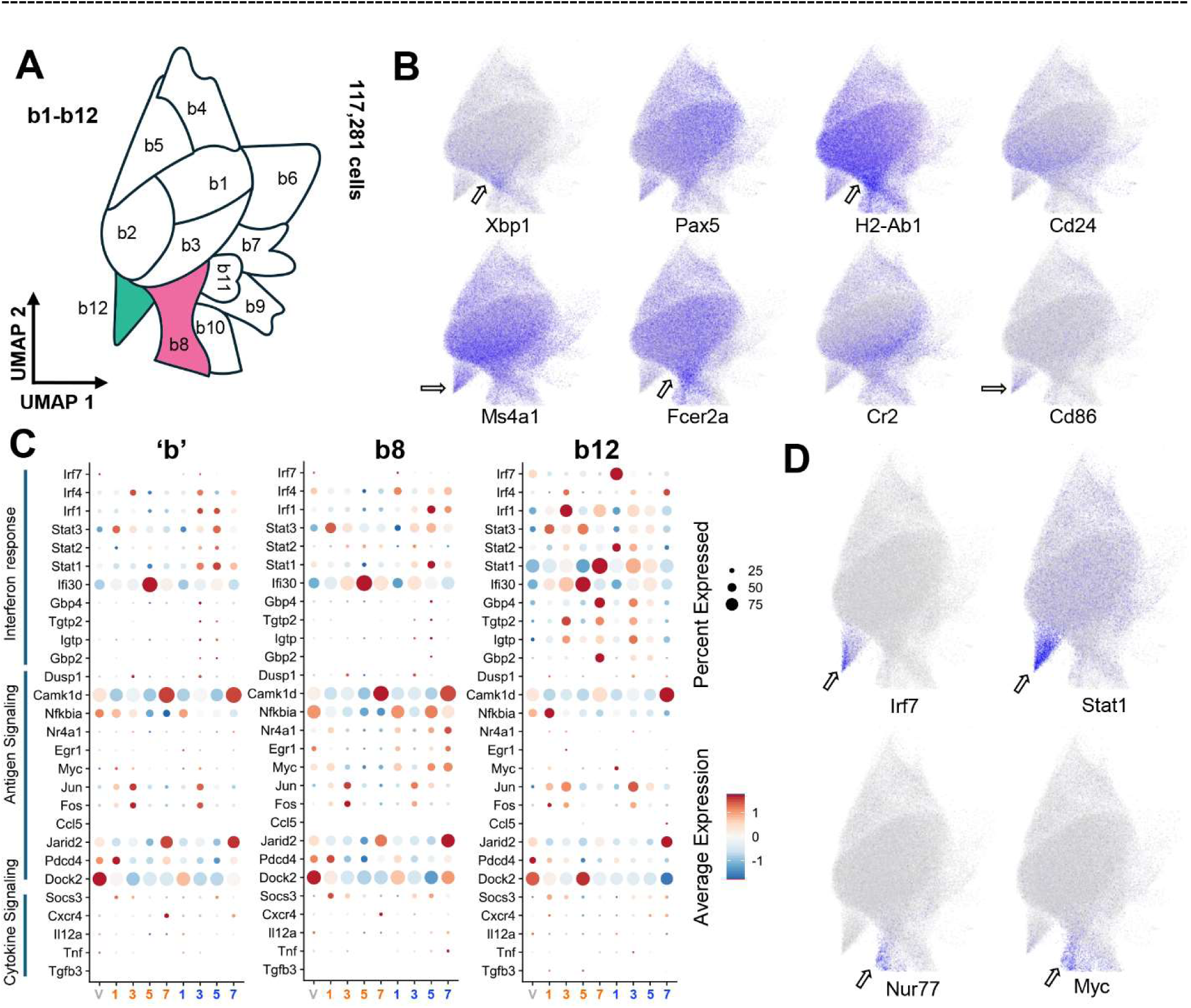
Exemplar dissection of the early behavior of specific B cell clusters following YF17D vs InYF immunization: (A) Representation of selected region of UMAP from Fig 1B for analysis of B cell clusters. b8 and b12 are further analyzed here. (B) Feature plots quantitating the expression of known B cell activation/differentiation markers Xbp1, Pax5, H2-Ab1 (MHC II), Cd24, Ms4a1 (Cd20), Fcer2a (Cd23), Cr2 (Cd21) and Cd86 in dLN. (Expression scale: <20^th^ percentile = gray, >80^th^ percentile = blue, 20-80 percentile → linearly scaled) (C) Key discriminating genes from b8 and b12 are compared to cells in the remaining B cell clusters (‘b’) across all timepoints (X-axis), color coded for VIPEX (V, gray), InYF (orange), YF17D (blue) respectively. Pseudobulk expressions for specific set of cells (b8, b12 or ‘b’) across timepoints are calculated and then scaled. The scale is saturated at 1.75 and -1.75 for consistent visualization across three plots. Cells with count for a given gene > 0 are included in the percentage expression calculation. (D) Unique markers discriminating b8 and b12: Nur77 (Nr4a1), Myc, Irf7 and Stat1.

Functionally, the b12 cluster was unique in that it was the sole IFN-responsive population within the B cell super-cluster (Figure S8A) as highlighted by the distribution of IRF7 and STAT1 transcripts (Figure 7D). It expressed a series of viral response genes elicited by YF17D (Figure S8A, 5A ‘LNXIII’, 7C ‘b12’), unlike any other B cell cluster. An expansion of these cells was also detected (Figure 3B, panel ‘b12, D3’) only in response to YF17D, specifically on D3. A kinetic analysis of key markers (Fig 7C) further illustrates this dichotomy. Intriguingly, Stat1 was activated on D7 in InYF, but on D3 in YF17D (Figure 7C, ‘b12’).

In contrast, cells in b8 display strong evidence of antigen receptor signaling (Figure 7B, 7C ‘b8’, S8B). This includes the upregulation of MHC II, Xbp1, Jun, Fos, IκBα (Figure 7C, ‘b8’ panel, S8B) while retaining a follicular phenotype (Cd23 high). Importantly, this subset was more dominant in response to InYF, D3 onwards (Figure 3B, ‘b8, D5’ panel).

Consistent with greater evidence of recent antigen-receptor signaling in b8, we also observe higher levels of both Egr-1^56^, c-myc^57^ and the canonical “strength of signaling” marker Nur77^58^. However, we did not find evidence that these cells were actively cycling, as Ki67 and cycling expressions were not markedly different. This analysis demonstrates the ability to map precise cell subsets to a high functional resolution – sufficient to generate key hypotheses for experimental validation. One such hypothesis could relate to the balance between IFN-responsive b12 subsets and antigen-stimulated b8 cells in determining the differential outcomes of vaccination with YF17D vs InYF – with one promoting bone marrow homing LLPCs while the other drives strongly stimulated but short-lived plasma cells. As discussed earlier, the enriched kinetic patterns in b8 and b12, had distinct trends against YF17D and InYF (Figure 5A, LNVII, LNXIII). Indeed, the depth of the resource presented in this dataset allows for several such hypotheses to be derived for illuminating key molecular and cellular events at the early stages post immunization in these benchmark vaccine-pairs so widely used for reverse vaccinology.

## Discussion

Systems vaccinology efforts to uncover early markers of effective and durable vaccine response require high resolution molecular maps of cellular events^17,59^. Here we generate a comprehensive resource with ∼2TB of data, profiling the responses in immunized “dirty” mice to YF17D and InYF. The scope of the dataset integrating data from close to 700 thousand cells, from 4 tissue sites across 6 timepoints is comparable to other valuable resources such as the COVID-19 immune atlas^60^ and blood atlas^61^, or the cytokine dictionary^62^ which sample a similar number of cells in 1-3 tissues. In addition, independent evaluation of male and female mice allows us to document shared and divergent single cell phenotypes in the two sexes. For instance, in almost every kinetic pattern, there were differences between males and females in the kinetic features of gene expression, such as context of response (YF17D or InYF), peak time, peak amplitude etc. (Figure 5, S5E, S6), more prominently so in spleen and PBMC. Kinetic differences across sexes were subtle for many patterns (i.e., the shared kinetics, see Figure S5E), however there were striking divergence in some cases (Figure S6). In pattern SPVII, for example, 65 genes from b6, elicited a stronger peak in response to InYF in males, to YF17D in females – both on D3 (Figure S6A, S7B). Several of these b6 genes were related to cell death, such as, Atp2a3, Serinc3, Ppia (intrinsic apoptosis due to ER/oxidative stress), Fth1, Tmem123 (necrotic cell death). Notably, global gene regulation in response to vaccines in PBMC was more influenced by sex (Figure 4J, 5C, S6B), compared to the dLN layer (Figure 4I, 5A, S5E). Recall that our PCA indicated that tissues are the largest source of variation in our sc-data (Figure 1C), which corroborated a few recent studies^63-66^. The tissue-associated nuances of sexual dimorphism in immune response highlighted here are relevant to appropriate vaccine designs relevant for the whole population.

In addition to the use of CITEseq affording a single cell resolution, a significant difference from previous studies on YF itself^34,35,39^ is the use of the VIPEX model. The choice of influenza, followed by Malaria and then a drug cure (Methods), while one of many possible permutations, is quite relevant to the prevalence of infectious diseases in many parts of the world, where advances in vaccine efficacy could have the highest impact on global health^28^.These are non-trivial perturbations to the basal state of SPF mice – flu affecting lung physiology^67^ while plasmodium induces extensive reorganization of multiple lymphocyte populations^68^. The inclusion of VIPEX samples (together with naïve SPF controls) therefore allows for an additional layer of enquiry. In our preliminary analysis we observed specific activated T-cell subsets (e.g., τ5 - Ccl5, Kdm5d) and metabolic reorganization of macrophages (m1 - Cd24a, Cox5a, Vcp) in PBMC in VIPEX, in contrast with naïve. A specific B cell subset, for instance, elicited enhanced expression of IL-12 and IL-6 pathway members (b6 - Laptm5, Irf8, Syk) in both PBMC and spleen. Certain B cell clusters (e.g., b4 – Cd74, H2-Aa) in lymph nodes had increased signatures of antigen presentation and processing (manuscript under preparation). Accordingly, while previous studies have the tacit complexity that the observed responses could be stemming from first inflammatory responses in a previously naïve host immune system, we expect these data to reflect specific differences resulting from YF or InYF.

The examination of gene expression within the 45 cell clusters also allows us to segregate three categories of markers for further inquiries (Figure 4). Many STOIC markers we report here should be potentially verifiable using bulk experiments (e.g., IRFs, STATs etc., which are already reported through previous resources^12,17,34,39^). FLAKEY markers, which require single-cell resolution for their discovery, can be a trove of important, unexplored vaccination markers. In some cases, this resolution helps map known markers to specific clusters - for instance, in PBMC, ♀, D3, Cd86 is upregulated in b1, b6 and b7 against YF17D but in d1 against InYF. Common markers, in contrast, are likely to be signatures of vaccination alone, independent of durability. Cxcr3, as an example, is upregulated in the latter context, in b6, against both vaccines. Notably, in relatively rare instances we observed certain genes which were upregulated in YF17D but downregulated in InYF, or vice-versa, in the same cluster or otherwise (Table S4, Figure S5A, ‘lead’ and ‘anti’ rows). Cd52, for example, is downregulated in the latter context, against YF17D in τ2, but upregulated in b6 against InYF.

The biological significance of the IFN responsive maturing B cell subset has not been previously reported. We could speculate that these represent very early indicators of LLPCs that eventually find residency in bone marrow niches. We did not evaluate bone marrow niches specifically since the timepoints focused on very early days, prior to significant seeding in that tissue. The other cluster we discuss, b8, represents cells with heightened antigen signaling response to the InYF formulation. The data is amenable to further segmentation. For instance, b8 may have interesting subpopulations – indicated by nuanced patterns of non-overlapping expression of markers such as Cr2, H2-Ab1 and Myc (Figure 7B, D). Another intriguing point is that we did not observe increased transcripts of IFNAR receptors in b12, raising questions of the timing and context of type I IFN signaling in these contexts.

Overall, our analysis validates the depth of the dataset generated here to compare the live and inactivated-adjuvanted Yellow Fever vaccines. Definition of STOIC and FLAKEY genes across all timepoints and tissues allows us to define new markers and pathways not apparent in previous studies. Superimposing the gene expression programs in certain cell clusters which formed coherent kinetic patterns and directed networks over time, together with known transcription factor motifs offers opportunities for additional modeling. With emerging profiles on multiple human vaccines, this resource can serve as a reference to relate parallel trends in peripheral blood to systems level dynamics.

## Supporting information

Supplementary Material (Chatterjee et al)

Supplementary Tables (Chatterjee et al)

## Acknowledgement

The project was funded by the Defense Advanced Research Projects Agency (DARPA) Assessing Immune Memory (AIM) program thought grant AIM-FP-009 to the MEMSEAL team. We thank the Flow Cytometry Shared Service of the University of Maryland Marlene and Stewart Greenebaum Comprehensive Cancer Center for flow cytometry assistance

## Methods

### Overall experimental design (Figure 1)

In this study, each mouse was pre-primed to generate Virus Immune Parasite Exposed (VIPEX, see below and REF) mice (Figure 1A). The VIPEX mice were then challenged with YF17D or inactive YF17D. To collect the single cell data analyzed in this report, groups of 6 mice (3 male (M) and 3 female (F)) are sampled on days 1, 3, 5, 7 and 30 after this challenge (Figure S1A-C). In addition, 2 (1 M, 1 F) VIPEX mice that were left unvaccinated are also analyzed at each of these days, as internal controls. Tissues are isolated and processed for CITEseq as discussed below.

For the design on D0 (Fig S1B) and post-D0 (Figure S1C), the samples of the same tissue from individual mice received hashtags^69^ and were pooled together before the cells were profiled using a droplet based system^70^ (10x Genomics, see Methods). High-quality sequence data of the single-cell transcriptome and cell-surface markers were generated (Illumina, see Methods) using the CITE-seq pipeline^71^.

### Generation of virus infected plasmodium-exposed (VIPEX) Mice

For the pre-vaccination (Day 0) timepoint, six male and six female C57BL/6 mice (The Jackson Laboratory) were used. Of these, half were unexposed naïve mice and half were sequentially exposed to two pathogens (VIPEX, see below for details). For each post-vaccination timepoint, seven male and seven female C57BL/6 mice (The Jackson Laboratory) were used, all VIPEX. Three of each gender were vaccinated either attenuated YF17D or inactivated YF17D. We also included un-vaccinated male and female control per each timepoint.

For VIPEX, 5-week-old mice were first infected intranasally (i.n.) with 10^5^ plaque forming units (pfu) of Influenza A/Puerto Rico/8/34 (PR8) virus. Mice were monitored by measuring body weight and allowed to recover for ∼2 weeks. After recovery from flu infection, mice were injected intraperitoneally (i.p.) with 10^5^ red blood cells (RBC) infected with *Plasmodium yoelii* strain 17XNL (BEI Resources). Disease progress was monitored by counting parasitemia in Giemsa-stained blood smears. Four doses of 1mg chloroquine were administered intraperitoneally over the course of 3 weeks to clear the malaria infection.

### Vaccination Regimen

Pre-exposed mice (now 12-15 weeks old) were injected intraperitoneally (i.p.) with 3mg of anti-mouse Interferon receptor alpha 1 (IFNAR-1) to prevent rapid clearance of the vaccine’s virus strains by type I IFN responses^72^. One day later (Day 0), mice were vaccinated subcutaneously (s.c.) in one caudal thigh muscle with live-attenuated virus yellow fever vaccine, YF17D (Sanofi Pasteur), or the formalin-inactivated YF17D.

Spleen, whole blood, draining lymph nodes (the ipsilateral inguinal and popliteal LN), and non-draining lymph nodes (the contralateral inguinal and popliteal LN) were harvested at the following timepoints: Day 0 (pre-vaccination), and Days 1, 3, 5, 7, and 30 post-vaccinations. Mice used for all timepoints later than Day 3 received additional i.p. injections of 0.6mg anti-mouse IFNAR-1 on Days 2 and 3 post-vaccination.

### Preparation of single-cell suspension

At each timepoint, whole blood was collected via cardiac puncture from anesthetized mice into tubes containing K_2_ EDTA anticoagulant (Greiner Bio-One). The whole blood was centrifuged at 2,000 rpm, after which the serum was removed. Red blood cells were lysed using Ack lysing buffer (Thermo Fisher Scientific) and peripheral blood mononuclear cells (PBMCs) were washed with Phosphate Buffered Saline (PBS) supplemented with 5% Fetal Bovine Serum (FBS).

Following blood collection, the spleen, draining LN, and non-draining LN were isolated from each mouse. Tissues were manually dissociated in PBS supplemented with 5% FBS and filtered through 70 μm nylon mesh.

From all tissues, live cells were isolated via density gradient centrifugation with Lympholyte M solution (Cedarlane Labs) and washed with PBS supplemented with 5% FBS. Cells were counted and viability was checked using acridine orange/propidium iodide (AOPI) staining solution (Nexcelom) and a Cellaca MX cell counter (Nexcelom).

### CITE-seq Staining

150,000 cells per sample were incubated with anti-mouse CD16/32 antibody (Trustain FcX Plus: BioLegend) to block non-specific binding to Fc receptors. Cells were then hashtagged using TotalSeq-C oligonucleotide-conjugated anti-mouse antibodies (BioLegend). Hashtagged cells were pooled together according to tissue type, and then stained with seventy-four TotalSeq-C cell lineage-specific anti-mouse surface proteins at concentrations ranging from 12.5ng to 100ng per pooled sample. Stained cells were washed with Cell Staining Buffer (BioLegend). Cells were counted and viability was checked using AOPI and a Cellaca MX cell counter. Normal viability range was 85-95%.

### Single Cell RNA-seq Preparation

We targeted the maximum input of 20,000 CITE-seq stained cells for Chromium Next GEM single cell 5’HT reagent kits v2 (Dual Index) (10x Genomics). Following the user guide (CG000424, Rev C), we constructed gene expression and cell surface protein libraries. Both libraries were pooled and sequenced using NovaSeq 6000 following 10x Genomics’ protocol (Read 1-26 cycles; i7 index-10 cycles; i5 index-10 cycles; Read 2-90 cycles).

### Sequencing and data extraction

The samples were pooled into batches and the base call (BCL) files from the sequencing instrument were demultiplexed using the mkfastq function of the Cell Ranger software (10x genomics), which generated fastq files containing the reads of different indexes. These fastq files were then processed by utilizing the count function of Cell ranger, and the filtered feature matrix barcode was utilized for all downstream analysis.

### Alignment

The transcript and surface reads for each index were aligned using the count function of CellRanger using the default settings. The CellRanger output, i.e., filtered bar code matrices were analyzed with Seurat (ver 4.3.0). 28 indexes of draining Lymph node samples, 25 indexes of spleen samples, 22 indexes of PBMC samples and 12 indexes of nondraining lymph nodes, hence altogether 87 indexes were first cleaned, then demultiplexed and finally integrated. We started the data processing with a total of 1,145,808 cells.

### Data pre-processing

Data processing was done in R (ver 4.3.0) relying largely on Seurat (ver 4.3.0). All cells which had more than 20,000 reads, had more than 2.5% of the reads coming from mitochondrial genes, or had less than 500 or more than 3000 unique features were removed from the dataset. Demultiplexing of pooled samples utilized HTODemux function (threshold = 0.999). Hashtags which did not meet this threshold were removed from our analysis.

The total number of samples in the experiments, each of which received one hashtag, divided for different 10x indexes were 1,130. After removing the correlated hashtags, and discarding the weakly stained hashtags (for whom, 0.999 percentile of lowest cluster was zero), followed by successful demultiplexing within each of the 87 indexes, 1,023 samples were included in the analysis. This quality control resulted in 870,061 cells remaining after cleanup and subsequently 666,410 cells remaining after demultiplexing. To remove potential confounding factors, TCR, BCR, ribosomal and mitochondrial genes were discarded before downstream analysis with Seurat.

Demultiplexed and processed indexes were normalized using SCtransform and integrated using the standard Seurat reciprocal PCA (RPCA) reduction workflow for finding the integration anchors. The data was then projected onto the first 50 principal components (RunPCA function, Seurat) with a resolution of 1.0 for finding neighbors and cell clusters using the Louvain algorithm (FindNeighbors, FindClusters functions, Seurat). We used Manhattan distances as the distance metric for the UMAP projection (RunUMAP function, Seurat). We fixed the randomness seeds at each step of the clustering and projection, for the purpose of reproducibility. This analysis led us to find 45 cell clusters in our data. Singletons were discarded before the further downstream analysis with 666,333 cells.

### Cataloguing the markers

Next, we used the findmarkers function in Seurat to search the cell surface and gene expression markers across all clusters 45 clusters. For cell surface markers we used the following cut offs: at least 50% cells in the cluster and 0.25-fold difference in binary log scale. For the gene expression markers, we used a more stringent cut off as follows: at least 50% cells in the cluster and 1.0-fold difference in the binary log scale. We catalogued all markers for both cell surface and gene expression for all 45 clusters in Table S1.

### Cell annotation

We took an approach of assigning broad and general identities to the clusters, without attempting to annotate them with refined immune cells identities. For instance, we wanted to know which clusters were B cells, but within the B cell clusters we did not attempt to assign which cluster is, for example, the marginal zone B cells. We observed that such attempts will make us merge more than one numeric cluster in many cases, and hence would lead to loss of resolution. We were more interested to see whether any gene expression differences between the YF17D and InYF vaccine were discoverable with a regular numerical clustering algorithm, without losing any resolution. Broad cell-type classifications were assigned based upon canonical cell-surface markers (see Table S1). Cell clusters were annotated with alphanumeric characters, B cells as b, T cell as τ, DCs as d, macrophages as m and so on (Fig 1B).

### Principal component analysis

We calculated aggregate expression for all cells, only T cells, only B cells and cells which are not lymphocytes. We used the RunPCA function from Seurat to calculate the principal components of surface expression (ADT), 3000 highly variable genes across tissues (HVG) and normalized RNA expression (SCtransformed, SCT) datasets.

### Differential gene expression analysis

We calculated the ‘pseudo-bulk’ expression^73^ by summing the read counts from each cell cluster, from each index and from each hashtag. We calculated the pseudo-bulk expression of each gene in each cell cluster from bioreplicates. Genes which were expressed in less than 5 cells among the total of 666,333 cells were removed from the analysis. Pseudo-bulk calculations were done independently in each tissue. Any cell cluster which had less than 50 cells for a given sex, a given vaccination type, and given timepoint (beyond D0), or any cell cluster with less than 50 cells for a given sex in unvaccinated exposed mice, were not considered in differential gene expression analysis.

After ‘pseudo-bulk’ing of the data we carried out differential gene expression analysis with utilizing the DESeq2 algorithm^41^. P-values from DESeq2 analyses were corrected with Bonferroni family-wise error rate correction method. Any gene in a cluster, with more than 0.25-fold (binary logarithm scale) difference and adjusted p-value below 0.05 were accepted as differential expression^62^.

### Variance decomposition analysis

We executed variance decomposition on the longitudinal data from each tissue separately, using the PALMO package in R^43^. We used sex, time, vaccine, clusters as the simple covariates and cluster-vaccine, cluster-vaccine-sex and cluster-vaccine-time as the conjugate covariates. Here the assumption is that the total variance in the expression of a gene is the sum of variation contributed from different experimental covariates and its residual variance. The contribution of a covariate was calculated as the proportion of total variance to that of the corresponding covariate. We used the normalized (SCtransformed) gene expression for these calculations. We restricted our calculations for the genes whose longitudinal average normalized gene expression > 0.2 (log-normalized RNA). We excluded timepoints D5 and D7 in this analysis to manage computational load of these very large calculations.

### Analysis of cell number dynamics

We calculated the total number of cells for a given time-sex-vax entry, separately for each tissue (supplementary file 3). In each tissue we removed the cell clusters which constituted of less than 0.1% of all cells from that tissue. Next, we normalized the cell counts from each cluster with the total number of cells in that specific time-sex-vaccine entry, i.e., we carried out a row-wise normalization, and column-wise standardization of the cell clusters (see Figure 3A, C, Table S1). The number of cells from separate mice and separate indexes from post-vaccination timepoints were compared with those of unvaccinated mice using two tailed Wilcoxon rank sum test, followed by Bonferroni correction to control family wise error rate.

### Identification of kinetic patterns

For each tissue, we picked up all differentially expressed gene in clusters, from all timepoints, both sexes and vaccines. Then, we calculated the average expression from the ‘pseudobulk’ed replicates for each of the aforementioned gene in clusters, for all timepoints, sex, and vaccination type combinations. From these we calculated ‘shrunken’ log fold changes^42^ across time-sex-vaccine type, compared to the sex-specific VIPEX, using DESeq2 package in R. Total number of unique gene in clusters for dLN, spleen and PBMC were 7559, 1847, and 2377 respectively.

Next, we attempted to find an optimal number of patterns to be searched in the data from each tissue. To that end, we calculated the Calinski-Harabsz (C-H) index for a series of potential number of patterns^74^ (Figure S5D). The number of patterns, beyond 10, where the maxima of C-H index was detected, was used further (Figure S5D). Visual inspection of hierarchically clustered patterns in each tissue indicated presence of at least more than 10 patterns in the data (Figure S5C). Any maxima in C-H index occurring for number of patterns below 10 were thus ignored.

We utilized partition around medoid^46^ (PAM) algorithm for searching the kinetic patterns, with number of allowed patterns fixed to the numbers stated previously (Figure S7X), utilizing the Cluster package in R^75^. We used PAM algorithm for finding medoids (i.e., exemplars) of the kinetic patterns^76^. After discarding the singletons, 30, 26 and 20 patterns were considered in dLN, spleen and PBMC datasets, respectively (Figure S5D, S6A, B), for down-stream analysis. Further, we used Fisher’s exact tests followed by Bonferroni correction for discovering the associations between kinetic patterns and cell-clusters. Very strong associations between certain kinetic patterns and specific cell clusters reinforced the appropriateness of our purely numeric cell clustering and annotation approach.

### Learning Bayesian networks

We collected sex-agnostic DEG-cluster events for construction of Bayesian networks at D1, D3 and D5. Pseudo-bulk’ed expression information of DEG-cluster (i.e., the nodes) was used for correlation analyses within Grow-shrink algorithm^50^ in bnlearn package^77^ in R. Briefly, it is a constraint-based structure learning algorithm which estimates Bayesian networks based on conditional independence tests^78^. It detects Markov blankets for each node using Pearson’s linear correlation as the test of conditional independence, with a significance threshold of 0.05. Solitary nodes were pruned for simplified visualizations (Figure 6, S7D, E).

## Data availability

Raw sequence reads are available in NCBI under the bioproject accession PRJNA1188714. Processed read counts and normalized, integrated dataset objects along with an interactive user interface will be made openly available via GEO and Single Cell Portal, respectively, upon successful publication of the manuscript.

