## Supplementary Material (Chatterjee et al) for "Kinetic patterns of single cell gene expression discriminate between the murine cellular responses to live attenuated and inactivated Yellow Fever vaccines"

This file includes:

Supplementary figures:

Figure S1: The details of experimental design and quality control.

Figure S2: Surface marker distributions and exploring the effects of experimental covariates on abundance of cells in cell-clusters.

Figure S3: Extended principal component analyses and summarized results from flow cytometry experiments.

Figure S4: Cell number dynamics and exploring the effects of covariates on gene expression.

Figure S5: Set theoretic description of gene groups in PBMC, variance decomposition analysis of gene expression in spleen, and kinetic pattern search in dLN.

Figure S6: Kinetic pattern search in spleen and PBMC.

Figure S7: List of enriched cell-clusters within kinetic-patterns and extended Bayesian networks.

Figure S8: Gene expression heatmaps superimposed on UMAPs from dLN dataset.

Description of supplementary tables:

Table S1: List of cell surface and transcript markers across cell clusters.

Table S2: Number of cells in each cell-cluster, from different tissues, across timepoints, exposure status, vaccination status and sex.

Table S3: List of differentially expressed genes from clusters.

Table S4: Gene groups across timepoint, sex and tissues.

Table S5: Fold change data with kinetic pattern annotations across timepoint, sex and vaccination status for all DEG-cluster entries from responding tissues.

Table S6: Enriched motifs in the promoters of genes from enriched cell-clusters within kinetic-patterns.

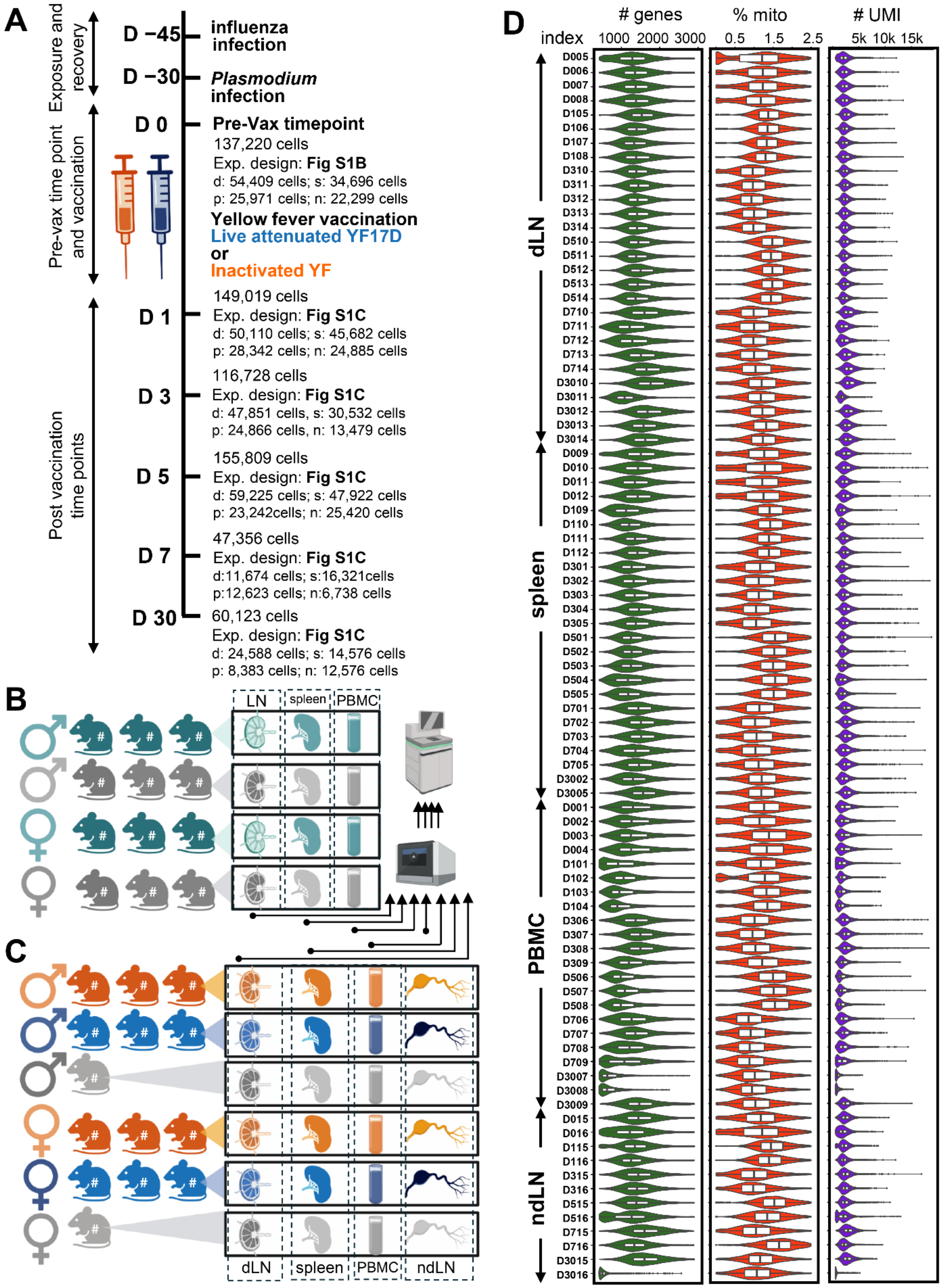


Fig S1: The details of experimental design and quality control.

(A) Cell numbers from each tissue at each time point is listed beside the experimental timeline (also see Table S2). Figures ‘S1B’ and ‘S1C’ is called across the timeline to refer to the corresponding experimental designs on specific timepoints. See below.

(B) The experimental design on pre-vaccination timepoint, i.e., Day 0. The ‘naïve’ cohort (dark green), the VIPEX cohort (dark grey) had male and female sub-cohorts (color-coded symbols on the left). The tissues shown in the callout boxes on the right (LN = lymph node, PBMC = peripheral blood mononuclear cells), were collected from each mouse in the cohort. The hash (#) sign on each mouse indicates that the tissues from each mouse received separate hashtags (see Methods for details). The vertical dashed boxes encompassing individual tissues indicate pooling of the individual tissue samples from all mice. These pooled samples were further run through Chromium 10x for single cell chemistry (solid-circle-tailed arrows) and then they were sequenced in Illumina NovaSeq (see Methods).

(C) The experimental design of each of the post-vaccination time points, i.e., Day 1, 3, 5, 7, and 30. Tissues were collected after specific time span after D0 from three cohorts of mice: cohort which received the live attenuated YF17D vaccine (blue), or inactivated YF vaccine (orange), or remained unvaccinated (light grey). Within these three cohorts there were sub-cohorts of male and female mice (color coded symbols on the left). Tissue samples, as shown in the callout boxes on the right, received hashtags representing individual mice. Figures (B) and (C) are made using BioRender.com.

(D) All indexes went through quality control filters. The percentage of mitochondrial genes, numbers of genes in a given cell and total number of reads from a given cell were filtered (see Methods). The distributions of the cells which were included in the analyses are shown here.

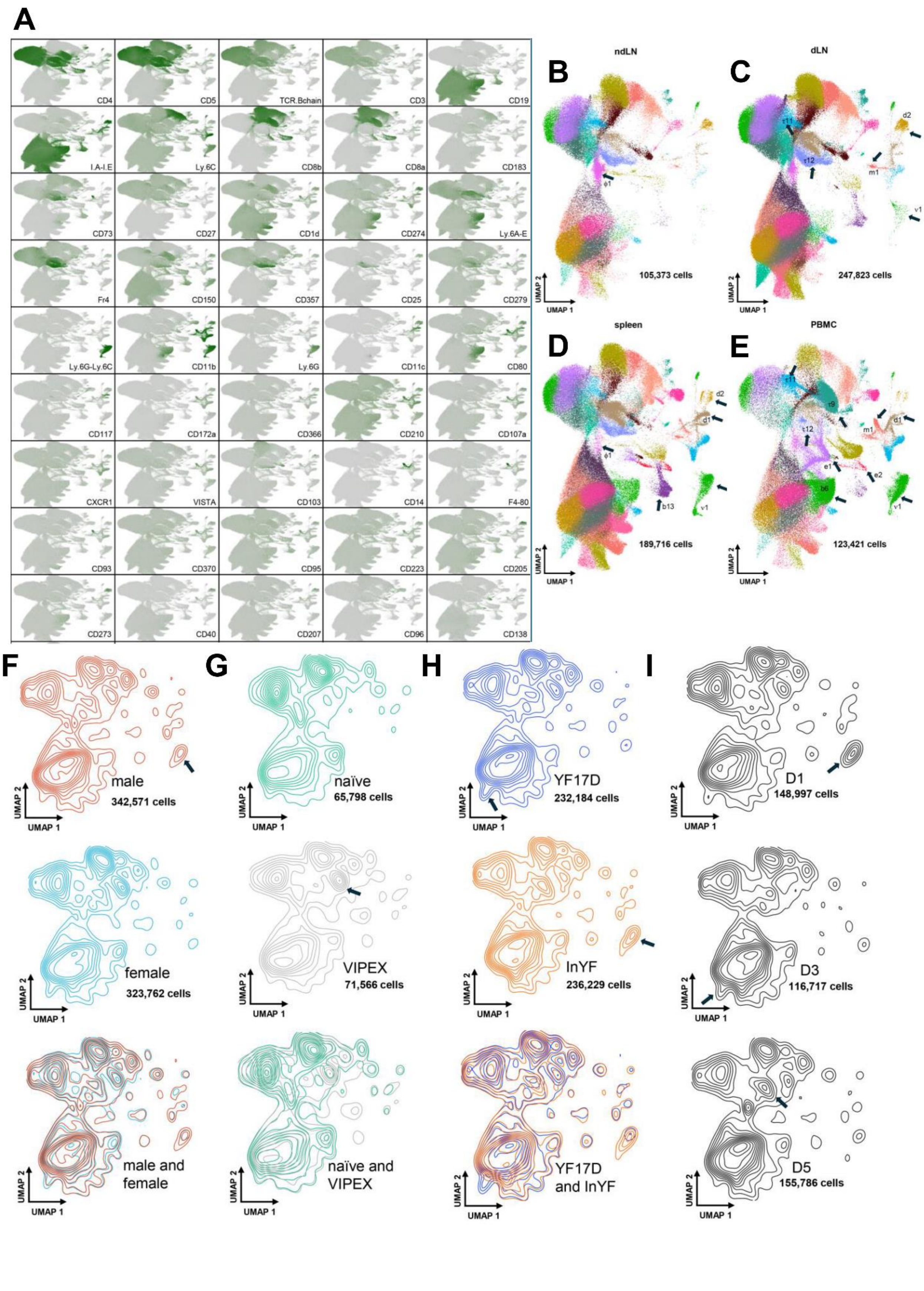
A
CD4
CD5
TCR.Bchain
CD3
CD19
I.A-I.E
Ly.6C
CD8b
CD8a
CD183
CD73
CD27
CD1d
CD274
Ly.6A-E
Fr4
CD150
CD357
CD25
CD279
Ly.6G-Ly.6C
CD11b
Ly.6G
CD11c
CD80
CD117
CD172a
CD366
CD210
CD107a
CXCR1
VISTA
CD103
CD14
F4-80
CD93
CD370
CD95
CD223
CD205
CD273
CD40
CD207
CD96
CD138
B
ndLN
105,373 cells
C
dLN
247,823 cells
D
spleen
189,716 cells
E
PBMC
123,421 cells
UMAP 1
UMAP 2
F
male
342,571 cells
female
323,762 cells
male and female
G
naïve
65,798 cells
VIPEX
71,566 cells
naïve and VIPEX
H
YF17D
232,184 cells
InYF
236,229 cells
YF17D and InYF
I
D1
148,997 cells
D3
116,717 cells
D5
155,786 cells

Fig S2: Surface marker distributions and exploring the effects of experimental covariates on abundance of cells in cell-clusters.

(A) Cell surface markers shown on the UMAP. (also see Table S1A).

(B-E) UMAPs for the whole dataset, showing the cells from individual tissues. Clusters where cell numbers markedly differ across tissues are marked with block arrows accompanied by the name of the cluster (see text). (B) ndLN, (C) dLN, (D) spleen, (E) PBMC.

(F-I) Contour maps of cell density for different experimental covariates. (F) Sex (male, female), (G) exposure status (naïve, VIPEX), (H) Vaccination status (YF17D, InYF), first two rows are superimposed in the third row. (I) time (D1, D3, D5). Block arrows in all figures indicate the differences discussed in the text.

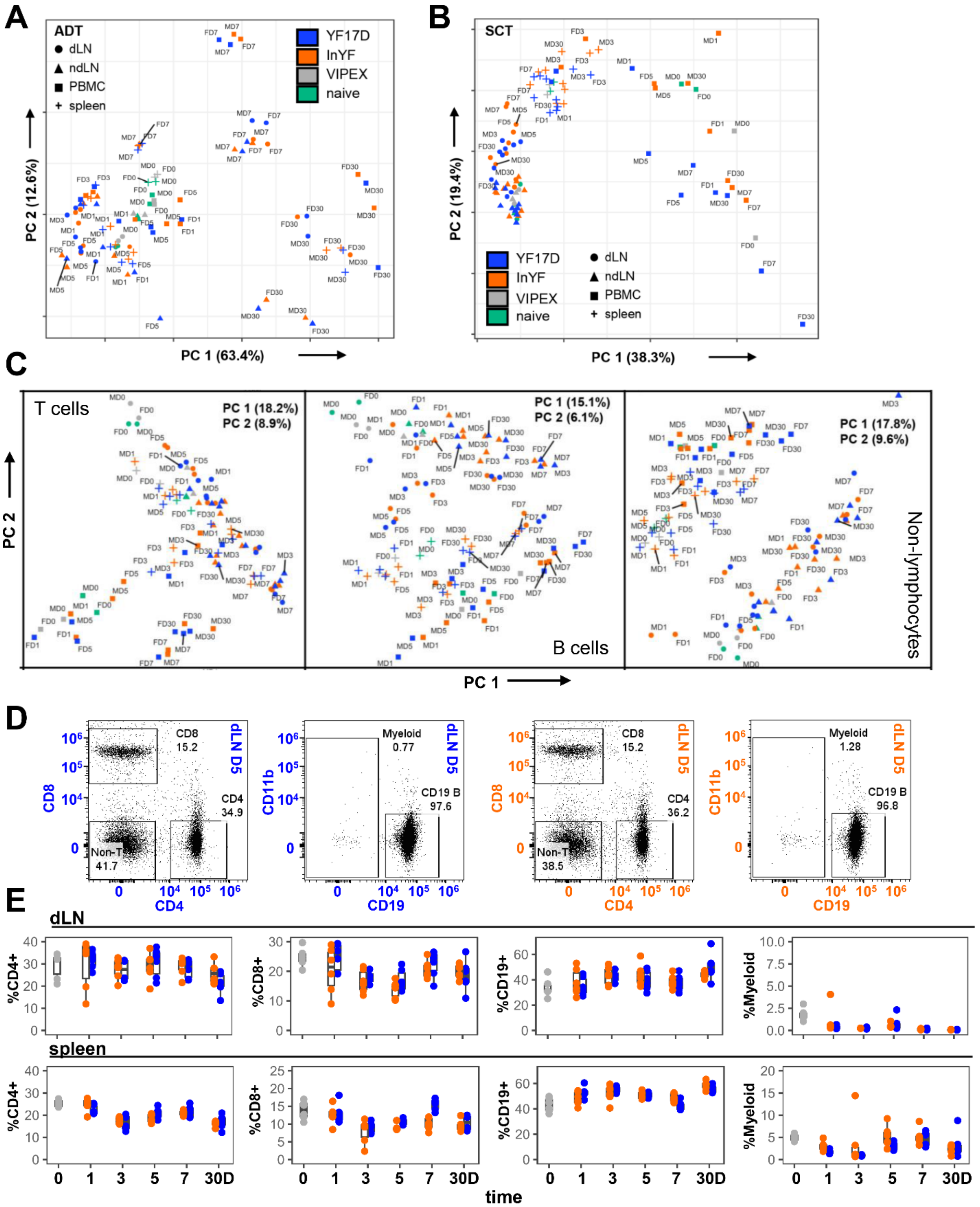


Fig S3: Extended principal component analyses and summarized results from flow cytometry experiments.

(A-C) Principal component analysis of bulk/semi-bulk version of the data. (A, B) Extension of figure 1C, PCA on ADT (antibody derived tag) and SCT (SCtransformed RNA) layers of whole dataset. (C) PCA on HVG (3000 highly variable genes across all data) layer from T cells, B cells and non-lymphocytes in the data. The legends for the plot symbols are same as in 1C, S3A, B.

(D) Representative flow cytometry data, depicting major immune markers CD4, CD8, CD19 and CD11b. The pair of scatter plots on the left, with blue title and axis labels represent the data from the dLN on D5 post-vaccination with YF17D. The orange ones are those from the mice vaccinated with InYF. In the CD4 vs CD8 scatters three gates are implemented: CD8+ T cells, CD4+ T cells, and Non-T cells. The latter population is separately analyzed in the CD19 vs CD11b scatters, where two gates are used: Myeloid cells and CD19+ B cells.

(E) Percentages of the CD8+ T cells, CD4+ T cells, CD19+ B cells and Myeloid cells were calculated from the flow analysis for the unvaccinated, and mice vaccinated with YF17D or InYF from different tissues across all timepoints. Data from dLN and spleen are shown. Absolute percentages of CD19+ B cells and Myeloid cells were calculated from the percentage of cells in non-T cell gate. Gray: unvaccinated VIPEX, blue: YF17D, orange: InYF. X-axis represents discrete time in days, and y-axis represents percentages. Wilcoxon Rank sum tests were performed on unvaccinated and vaccinated datasets and Bonferroni correction was applied for multiple correction. None of the differences were statistically significant at 0.01 level.

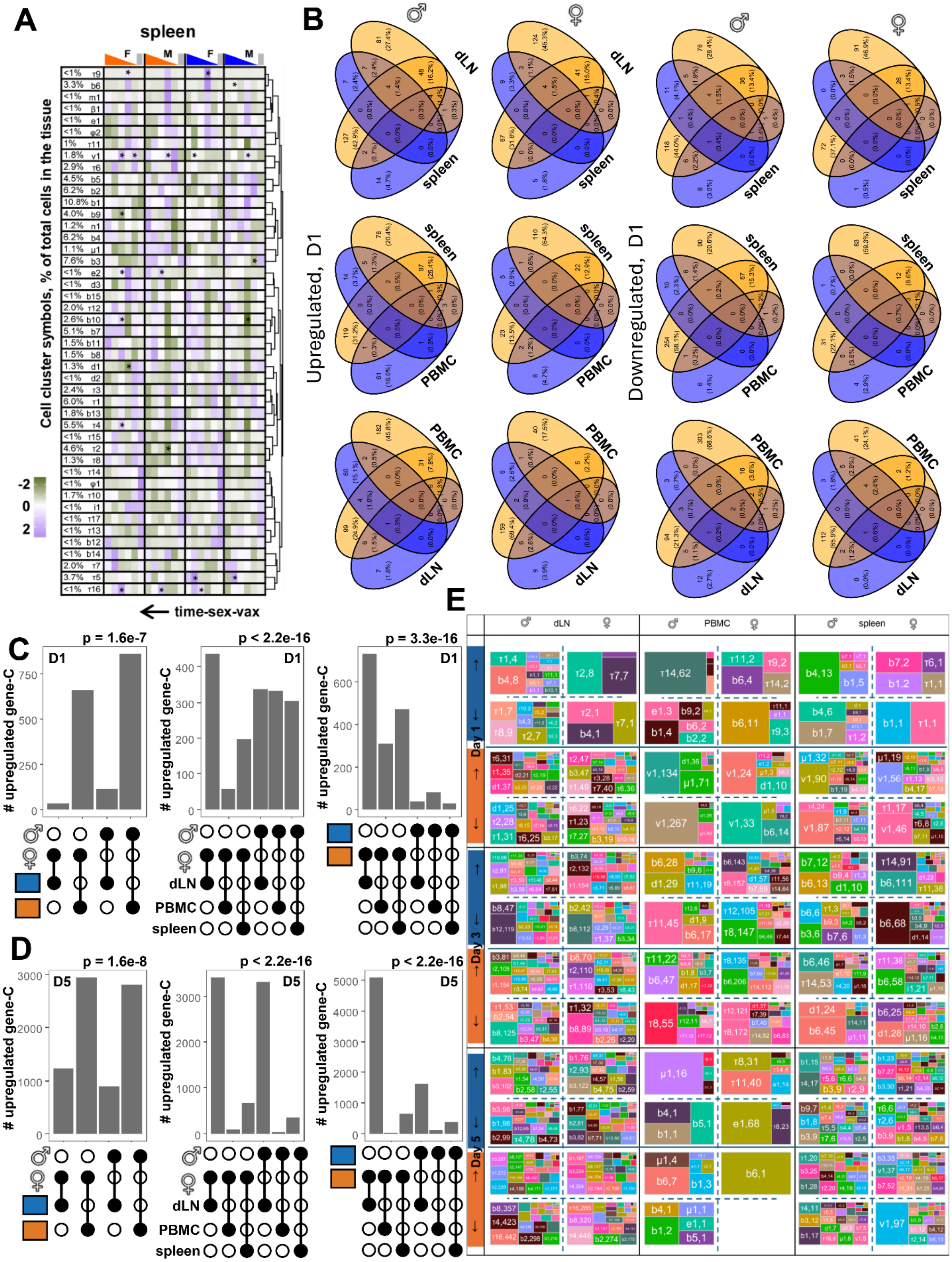


80 Fig S4: Cell number dynamics and exploring the effects of covariates on gene expression.

(A) Cell number dynamics in spleen. Description same as Figure 3A.

(B) Venn diagrams for exploring the overlap of gene expression across tissues. Data from upregulated D1, downregulated D1 are shown. Conclusion from other time points were similar (not shown due to space constraints).

(C-D) Up-set plots for exploring associations between experimental covariates, sex, vaccine and tissue on (C) D1, (D) D3, for the number of upregulated DEGs from clusters. P-values are from G-test of independence on the data shown in the plots. Conclusion from other time points, for both upregulated and downregulated DEG-cluster events were similar (not shown due to space constraints).

(E) Treemaps to show the source cell clusters of differential gene expression, across sex, vaccine, tissues, time points, and up regulation and down regulation. Colors are randomly picked and are unrelated to the colors in any other figure. Cluster symbols are printed in the rectangles, within treemaps, followed by the number of genes differentially expressed from that cluster.

**A** *Λ*: DEGs from InYF; *Γ*: DEGs from YF17D — **PBMC, D3 ♀**

| Gene group | Description | # | e.g. |
|---|---|---|---|
| common↑ | **(*Γ*↑-C)∩(*Λ*↑-C)** | 497 | Dusp2-τ1, Jun-τ1, Ier2/τ1, Junb/τ1, Tnfaip3/τ1 |
| common↓ | **(*Γ*↓-C)∩(*Λ*↓-C)** | 316 | Pip4k2a-τ1, Serf2-b1, Crip1-b1, Vim-b3 |
| STOIC, g↑, b↑, g↓, b↓ | ***Γ*↑ −*Λ*↑,*Γ*↑ −*Λ*↑, *Γ*↓ −*Λ*↓,*Γ*↓ −*Λ*↓** | 148, 236, 83, 256 | Snx5-b1, Ifrd1-b1, Ifrd1-d1, Ubb-b2, Gem-b3 B3gnt7-b6, Cxcr4-b6, Slamf1-b6, Hsd11b1-b6 Ass1-b6, Acot7-b6, Tbcb-b6, Nme1-b6, Crip2-b6 Myl6-τ5, Ctla4-b7, Pard3b-b7, Pkig-b7, Cd2-b7 |
| FLAKY, g↑, b↑, g↓, b↓ | **(*Γ*↑∩*Λ*↑) − common↑ (*Γ*↓∩*Λ*↓) − common↓** | 175, 153, 93, 148 | Nsa2-τ1, Cd86-b1, Lncpint-τ2, Stat4-b6, Cd55-b6 Nsa2-τ8, Cd86-d1, Lncpint-τ11, Stat4-τ8, Cd55-i1 Airn-τ2, Lgals1-b6, S100a13-b7, Ost4-b7, Ccnd2-b7 Airn-τ14, Lgals1-τ2, S100a13-d1, Ost4-b6, Ccnd2-b6 |
| common lead | **(*Γ*↓-C)∩(*Λ*↑-C)** | 0, 0 | - |
| common anti | **(*Γ*↑-C)∩(*Λ*↓-C)** | 0, 0 | - |
| general lead, g↓, b↑ | **(*Γ*↓∩*Λ*↑) − common lead** | 44, 38 | Cd247-τ8, Rnf157-τ8, Myc-τ12, Sytl3-τ12 Cd247-i1, Rnf157-b6, Myc-τ11, Sytl3-b6 |
| general anti, g↑, b↓ | **(*Γ*↑∩*Λ*↓) − common anti** | 50, 56 | Btg2-τ1, Jund-b3, Ctla4-τ2, Cebpb-τ2 Btg2-τ8, Jund-v1, Ctla4-b7, Cebpb-d1 |

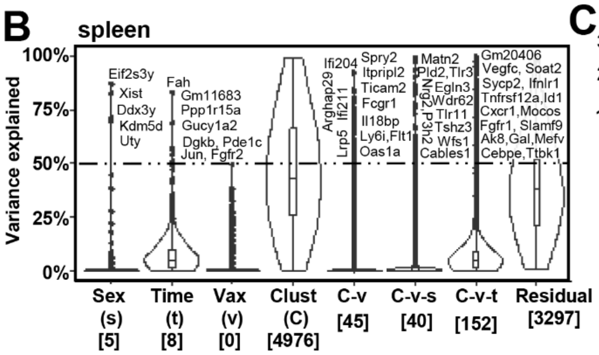


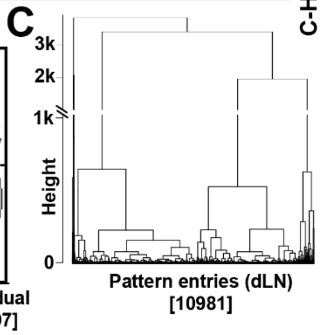


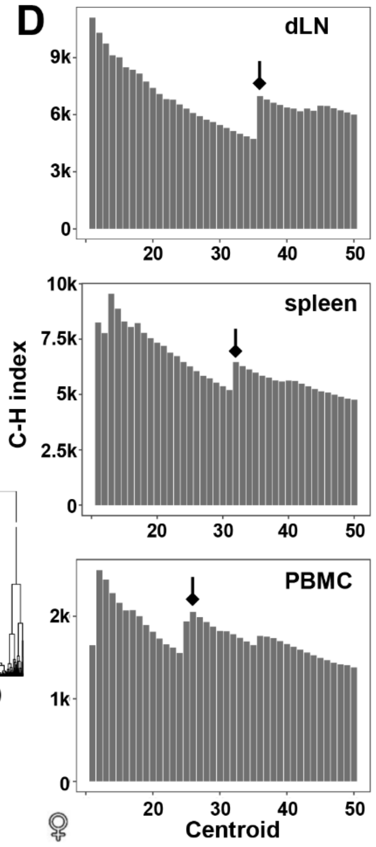


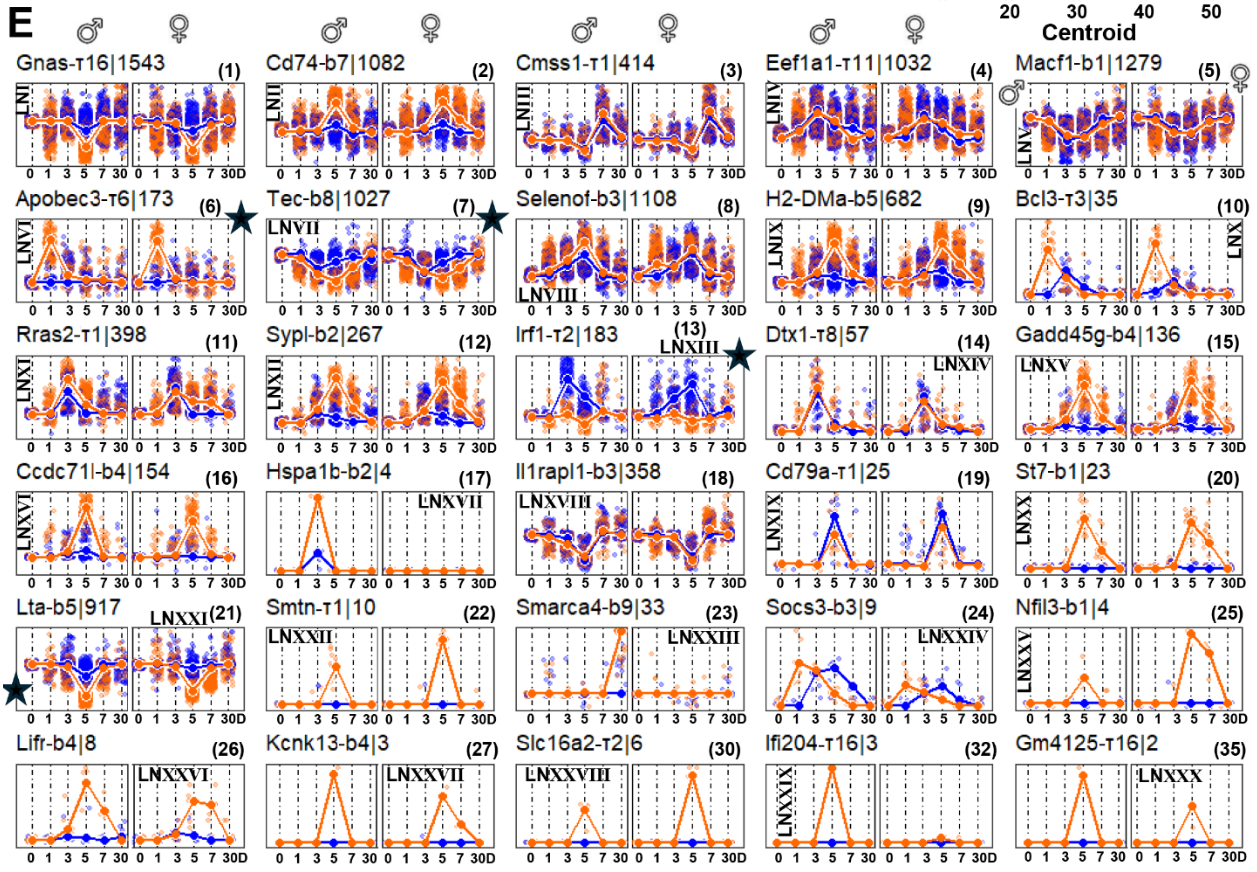

Fig S5: Set theoretic description of gene groups in PBMC, variance decomposition analysis of gene expression in spleen, and kinetic pattern search in dLN.

(A) Description complimentary to Figure 4A, B. Lists of gene groups, defined by set theoretic operations on differentially expressed gene distributions (from Figure 3D). Any DEG can be either of the three groups: common, STOIC or FLAKY. Adding common ($\times 2$), STOIC, FLAKY entries will match with total number of DEG-cluster events, for a given tissue, time, sex. The up arrow (↑) should be read as 'up': STOIC YF17D up, FLAKY InYF down etc. InYF is abbreviated as $\Lambda$ (lambda) and YF17D is abbreviated as $\Gamma$ (gamma), (mentioned at the top). **(Γ↑-C):** Upregulated in YF17D, from specific clusters. **(Γ↑ -Λ↑):** Set difference between genes upregulated in YF17D and upregulated genes in InYF, when cell cluster information is removed. In the description column all expressions are not provided for FLAKY groups. Only FLAKY YF17D up and down are shown. The 'lead' (downregulated in YF17D and upregulated in InYF) and 'anti' (upregulated in YF17D and downregulated in InYF) gene groups are shown at the bottom. The names 'lead' and 'anti' are borrowed from lead and anti-diagonals in geometry, to describe the diagonal relationships in gene expression (see Figure 4A to see the diagonal relations). Data from PBMC, D3, female. For list of gene groups for all tissues, timepoints and sex see Table S4.

(B) Variance decomposition analysis of gene expression data from spleen. Description same as Figures 4A, B. (C) Visual inspection of dendrograms of hierarchical clustering of all kinetic patterns from dLN. (D) Calinski-Harabasz indexes (C-H index) for all responding tissues for a range of centroid numbers. The second maxima beyond centroid = 15 were used for further analysis in each case (arrowheads).

(E) Kinetic patterns in dLN, log2FC > 0.25 or log2FC < -0.25, FWER adjusted p-value < 0.05. Pattern search with partition around medoid (PAM) algorithm with k = 36. In the title of each plot, the medoid gene-cluster and total number of gene-cluster entries in that pattern is denoted, separated by a vertical bar. Each subplot is basically a pair of two smaller plots: on the left data from male mice, on the right data from female mice. X-axis represents time: 0, 1, 3, 5, 7 and 30 days. Orange is data from the mice which received InYF and blue, from YF17D. Each plot has a cloud of orange and blue dots in the background. These are standardized fold changes from individual gene-cluster entries within a given pattern. In the foreground, the temporal pattern of the medoid gene-cluster is shown. Each subplot has a pattern id, 'LN' followed by roman numeric – these ids are used in main text. There are also corresponding serial numbers in the round brackets at top-right in each plot. These exact numbers are used in Table S5A, column 'pattern', for convenient machine-readability. There were six singleton patterns, which contained the following gene-cluster entries respectively, and which were not pursued further: Nfil3-b3, Efna5-τ2, Nfil3-b7, Nfil3-b8, Efna5-τ10, Lgals7-τ16. dLN, Bonferroni FWER correction, log2FC > 0.25 or log2FC < -0.25 dataset had a total of 10,981 gene-cluster entries which were explored for the kinetic pattern search. Plots with large star-shaped symbols are the ones discussed in the main text.

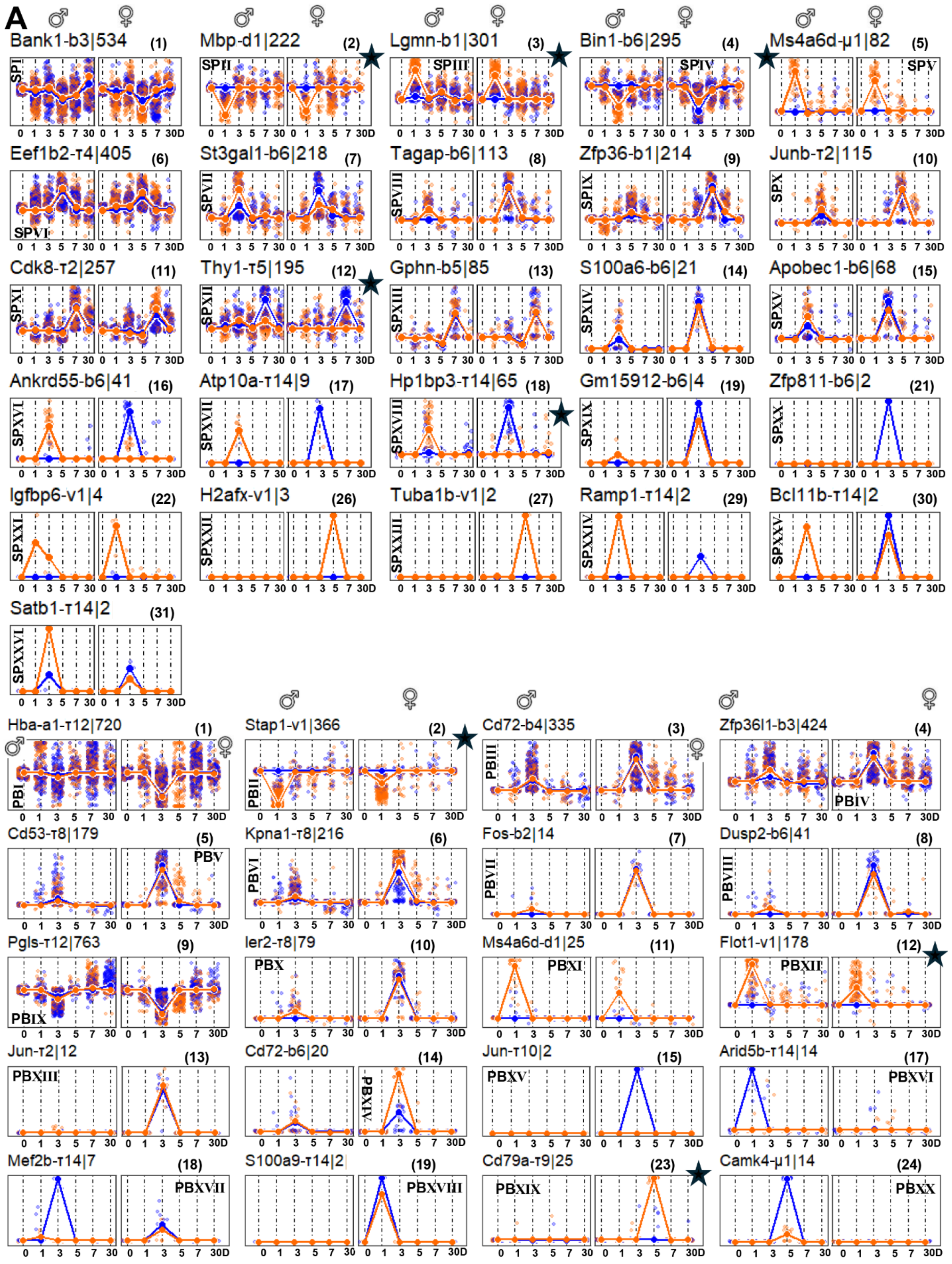
A
Bank1-b3|534 (1)
Mbp-d1|222 (2)
Lgmn-b1|301 (3)
Bin1-b6|295 (4)
Ms4a6d-µ1|82 (5)
Eef1b2-τ4|405 (6)
St3gal1-b6|218 (7)
Tagap-b6|113 (8)
Zfp36-b1|214 (9)
Junb-τ2|115 (10)
Cdk8-τ2|257 (11)
Thy1-τ5|195 (12)
Gphn-b5|85 (13)
S100a6-b6|21 (14)
Apobec1-b6|68 (15)
Ankrd55-b6|41 (16)
Atp10a-τ14|9 (17)
Hp1bp3-τ14|65 (18)
Gm15912-b6|4 (19)
Zfp811-b6|2 (21)
Igfbp6-v1|4 (22)
H2afx-v1|3 (26)
Tuba1b-v1|2 (27)
Ramp1-τ14|2 (29)
Bcl11b-τ14|2 (30)
Satb1-τ14|2 (31)
Hba-a1-τ12|720 (1)
Stap1-v1|366 (2)
Cd72-b4|335 (3)
Zfp36l1-b3|424 (4)
Cd53-τ8|179 (5)
Kpna1-τ8|216 (6)
Fos-b2|14 (7)
Dusp2-b6|41 (8)
Pgls-τ12|763 (9)
Ier2-τ8|79 (10)
Ms4a6d-d1|25 (11)
Flot1-v1|178 (12)
Jun-τ2|12 (13)
Cd72-b6|20 (14)
Jun-τ10|2 (15)
Arid5b-τ14|14 (17)
Mef2b-τ14|7 (18)
S100a9-τ14|2 (19)
Cd79a-τ9|25 (23)
Camk4-µ1|14 (24)
0 1 3 5 7 30 0 1 3 5 7 30D

Fig S6: Kinetic pattern search in spleen and PBMC. (A) spleen (B) PBMC. Description same as S5E. (A) Pattern ids: 'SP' followed by Roman numerals. Total number of gene-cluster entries: 3267. k = 32. Six singletons: Tppp3-b6, Prok2-v1, Nrgn-v1, Ctla2a-v1, S100a9-μ1, Sh2d1a-τ14. (B) Pattern ids: 'PB' followed by Roman numerals. Number of entries: 3442. k = 26. Six singletons: Fos-b9, Pik3ap1-τ14, Igfbp6-d1, Prok2-v1, Ppp1r15a-τ8, Fos-τ12.

**A**

| pattern | cluster | OR | % | #gene | padjust |
|---|---|---|---|---|---|
| 1 | b6 | 2.97 | 2.2 | 34 | 3.5573E-05 |
| 4 | τ1 | 2.78 | 13.57 | 140 | 3.3416E-19 |
| 4 | τ11 | 2.94 | 4.17 | 43 | 7.8705E-07 |
| 4 | τ13 | 3.01 | 2.91 | 30 | 5.88E-05 |
| 4 | τ17 | 5.39 | 0.97 | 10 | 0.00433898 |
| 5 | τ13 | 3.66 | 3.21 | 41 | 1.7172E-08 |
| 5 | b12 | 20.61 | 11.88 | 152 | 1.5884E-92 |
| 6 | τ6 | 4.4 | 9.83 | 17 | 4.638E-05 |
| 6 | b6 | 4.6 | 4.05 | 7 | 0.03466486 |
| 6 | τ13 | 4.92 | 5.2 | 9 | 0.0048528 |
| 6 | d1 | 37.21 | 15.03 | 26 | 6.0848E-27 |
| 6 | d2 | 10.92 | 8.67 | 15 | 3.1952E-09 |
| 7 | b8 | 11.33 | 32.42 | 333 | 2.666E-159 |
| 8 | b4 | 2.89 | 11.82 | 131 | 4.2283E-19 |
| 8 | τ7 | 3.02 | 6.59 | 73 | 8.1915E-12 |
| 9 | d2 | 3.07 | 2.64 | 18 | 0.00302747 |
| 10 | d1 | 46.52 | 22.86 | 8 | 1.3079E-09 |
| 11 | τ3 | 3.67 | 8.29 | 33 | 1.1295E-07 |
| 11 | τ6 | 3.39 | 7.54 | 30 | 2.5192E-06 |
| 11 | τ8 | 2.54 | 5.28 | 21 | 0.00806041 |
| 11 | τ15 | 7.76 | 2.51 | 10 | 0.00010784 |
| 11 | τ17 | 17.64 | 2.76 | 11 | 4.5962E-08 |
| 11 | b14 | 22.74 | 9.05 | 36 | 4.4217E-29 |
| 13 | τ2 | 2.88 | 15.85 | 29 | 0.0001017 |
| 13 | τ10 | 2.82 | 8.2 | 15 | 0.01888143 |
| 13 | τ12 | 3.19 | 8.74 | 16 | 0.00366511 |
| 13 | τ13 | 4.06 | 4.37 | 8 | 0.03743682 |
| 13 | b12 | 3.32 | 6.01 | 11 | 0.0264757 |
| 14 | τ17 | 45.42 | 8.77 | 5 | 6.8018E-06 |
| 14 | b14 | 36.45 | 19.3 | 11 | 6.7962E-12 |
| 15 | b10 | 3.27 | 7.35 | 10 | 0.04072274 |
| 16 | τ16 | 2.54 | 14.29 | 22 | 0.00645819 |
| 16 | d2 | 15.77 | 11.69 | 18 | 2.0754E-13 |
| 18 | b7 | 3.11 | 8.66 | 31 | 7.976E-06 |
| 18 | τ16 | 2.6 | 14.25 | 51 | 6.9085E-07 |
| 19 | τ1 | 7.3 | 32 | 8 | 0.00095747 |
| 19 | τ2 | 5.83 | 28 | 7 | 0.00814101 |
| 21 | τ4 | 4.65 | 27.59 | 253 | 1.4491E-63 |
| 21 | τ16 | 20.31 | 40.13 | 368 | 4.042E-233 |
| 23 | b9 | 214.33 | 66.67 | 22 | 2.9839E-35 |
| 24 | d1 | 41.37 | 22.22 | 2 | 0.0135539 |
| 27 | b4 | 36.66 | 66.67 | 2 | 0.01554597 |
| 32 | τ16 | 29.93 | 66.67 | 2 | 0.02262941 |

**B**

| Pattern | cluster | OR | % | #gene | padjust |
|---|---|---|---|---|---|
| 1 | τ5 | 2.66 | 12.17 | 65 | 1.3331E-07 |
| 2 | v1 | 18.55 | 58.11 | 129 | 9.594E-76 |
| 3 | b4 | 3.11 | 11.63 | 35 | 6.4595E-06 |
| 3 | μ1 | 2.97 | 10.63 | 32 | 4.4774E-05 |
| 4 | b6 | 8.53 | 38.31 | 113 | 9.9153E-45 |
| 4 | d1 | 6.66 | 16.61 | 49 | 8.2576E-18 |
| 5 | v1 | 16.42 | 62.2 | 51 | 4.0179E-30 |
| 5 | μ1 | 6.7 | 21.95 | 18 | 7.3982E-08 |
| 6 | b1 | 2.88 | 12.1 | 49 | 5.6624E-07 |
| 6 | b3 | 2.58 | 11.36 | 46 | 1.7887E-05 |
| 6 | τ2 | 2.61 | 8.64 | 35 | 0.00026739 |
| 7 | τ4 | 3.06 | 11.93 | 26 | 0.00020287 |
| 7 | b6 | 4.76 | 29.82 | 65 | 3.8967E-17 |
| 7 | e1 | 14.24 | 2.29 | 5 | 0.00682316 |
| 8 | b6 | 4.61 | 30.97 | 35 | 1.6118E-09 |
| 8 | τ11 | 25.39 | 27.43 | 31 | 1.2713E-25 |
| 8 | i1 | 10.3 | 5.31 | 6 | 0.00179556 |
| 9 | b7 | 3.14 | 14.49 | 31 | 2.3062E-05 |
| 10 | τ13 | 11.84 | 4.35 | 5 | 0.00704954 |
| 10 | τ17 | 23.7 | 4.35 | 5 | 0.00062415 |
| 10 | τ16 | 15.56 | 6.96 | 8 | 1.8917E-05 |
| 11 | b5 | 3.1 | 6.23 | 16 | 0.00867905 |
| 11 | b10 | 4.63 | 3.89 | 10 | 0.00846014 |
| 12 | τ5 | 11.17 | 33.85 | 66 | 9.7451E-34 |
| 13 | τ7 | 6.53 | 8.24 | 7 | 0.00616247 |
| 14 | b6 | 12.95 | 57.14 | 12 | 4.8135E-07 |
| 15 | b6 | 7.11 | 41.18 | 28 | 3.1188E-11 |
| 15 | τ14 | 5.08 | 17.65 | 12 | 0.00024181 |
| 15 | τ11 | 7 | 13.24 | 9 | 0.00022748 |
| 16 | τ14 | 52.22 | 65.85 | 27 | 1.9897E-27 |
| 17 | τ14 | 45.96 | 66.67 | 6 | 1.7766E-06 |
| 18 | d1 | 3.39 | 12.31 | 8 | 0.03369369 |
| 18 | τ14 | 78.44 | 70.77 | 46 | 3.0307E-50 |
| 22 | v1 | Inf | 100 | 4 | 0.00011631 |
| 26 | v1 | Inf | 100 | 3 | 0.00112446 |
| 27 | v1 | Inf | 100 | 2 | 0.01084202 |
| 29 | τ14 | Inf | 100 | 2 | 0.00185686 |
| 30 | τ14 | Inf | 100 | 2 | 0.00185686 |
| 31 | τ14 | Inf | 100 | 2 | 0.00185686 |

**D**

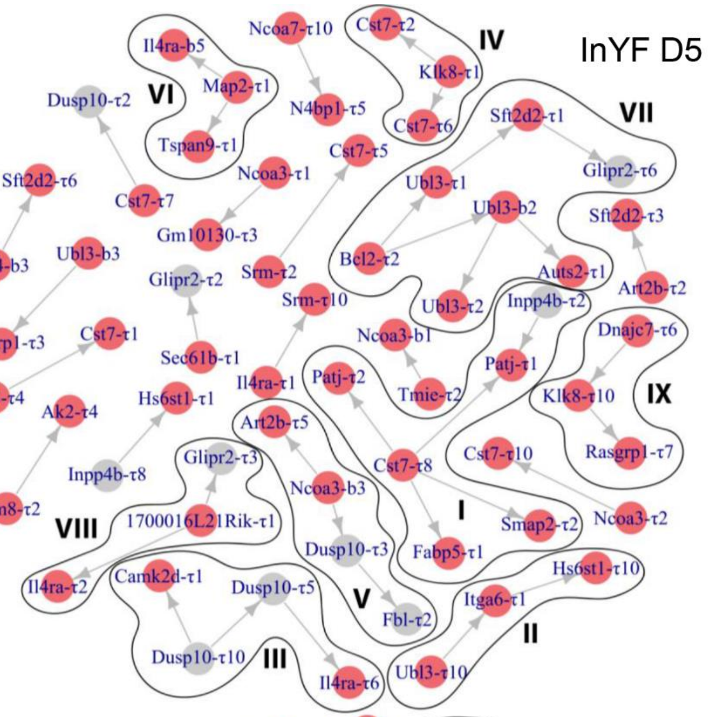


**E**

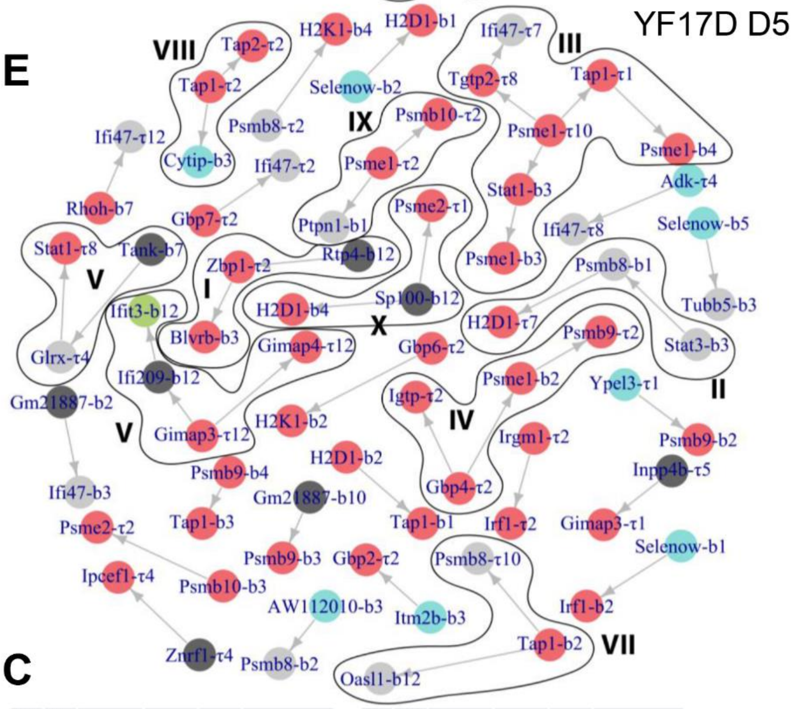


**C**

| pattern | cluster | OR | % | #gene | padjust |
|---|---|---|---|---|---|
| 1 | b1 | 3.91 | 4.86 | 35 | 1.2212E-06 |
| 1 | τ5 | 6.76 | 6.81 | 49 | 1.4492E-14 |
| 1 | e1 | 20.25 | 12.5 | 90 | 3.1535E-43 |
| 1 | τ9 | 31.15 | 11.25 | 81 | 3.436E-43 |
| 2 | v1 | 59.48 | 76.5 | 280 | 1.266E-212 |
| 2 | μ1 | 3.6 | 11.2 | 41 | 1.5188E-08 |
| 4 | b4 | 3.87 | 4.95 | 21 | 0.00013071 |
| 4 | b6 | 4.08 | 31.37 | 133 | 9.7051E-27 |
| 4 | b9 | 4.98 | 3.07 | 13 | 0.00116375 |
| 4 | τ1 | 3.78 | 5.66 | 24 | 4.0118E-05 |
| 5 | τ8 | 3.05 | 26.26 | 47 | 1.6819E-07 |
| 6 | τ8 | 2.97 | 25.46 | 55 | 2.1527E-08 |
| 6 | b7 | 2.63 | 8.8 | 19 | 0.00986223 |
| 7 | n1 | 183.49 | 14.29 | 2 | 0.00153127 |
| 8 | τ14 | 3.49 | 26.83 | 11 | 0.02021779 |

| pattern | cluster | OR | % | #gene | padjust |
|---|---|---|---|---|---|
| 8 | τ3 | 33.21 | 7.32 | 3 | 0.00365404 |
| 9 | τ8 | 4.43 | 25.56 | 195 | 1.3808E-37 |
| 9 | τ12 | 13.57 | 21.89 | 167 | 7.9407E-68 |
| 9 | τ7 | 19.05 | 6.68 | 51 | 2.5244E-23 |
| 11 | v1 | 12.54 | 64 | 16 | 1.5949E-08 |
| 12 | d1 | 2.84 | 14.61 | 26 | 0.00024793 |
| 12 | v1 | 8.74 | 51.12 | 91 | 1.2774E-36 |
| 12 | μ1 | 9.84 | 24.16 | 43 | 7.3933E-22 |
| 14 | τ14 | 6.31 | 40 | 8 | 0.0016214 |
| 15 | τ10 | 534.02 | 50 | 1 | 0.00814189 |
| 17 | τ14 | Inf | 100 | 14 | 5.2393E-15 |
| 18 | τ14 | Inf | 100 | 7 | 7.7451E-08 |
| 19 | τ14 | Inf | 100 | 2 | 0.00942347 |
| 23 | τ11 | 99.66 | 88 | 22 | 4.4753E-22 |
| 24 | μ1 | Inf | 100 | 14 | 3.0514E-20 |

Fig S7: List of enriched cell-clusters within kinetic-patterns and extended Bayesian networks.

(A) Association between clusters and patterns in dLN. One tailed Fisher’s exact tests were executed to find positive associations between clusters and patterns. P-values were further adjusted with FWER, i.e., Bonferroni approach. Odds ratios > 2.5, adjusted p-value < 0.05 are listed. OR – Odds ratio.

(B) Association between clusters and patterns in spleen.

(C) Association between clusters and patterns in PBMC.

(D, E) Bayesian network for InYF D5 and YF17D D5. Description same as Figure 6.

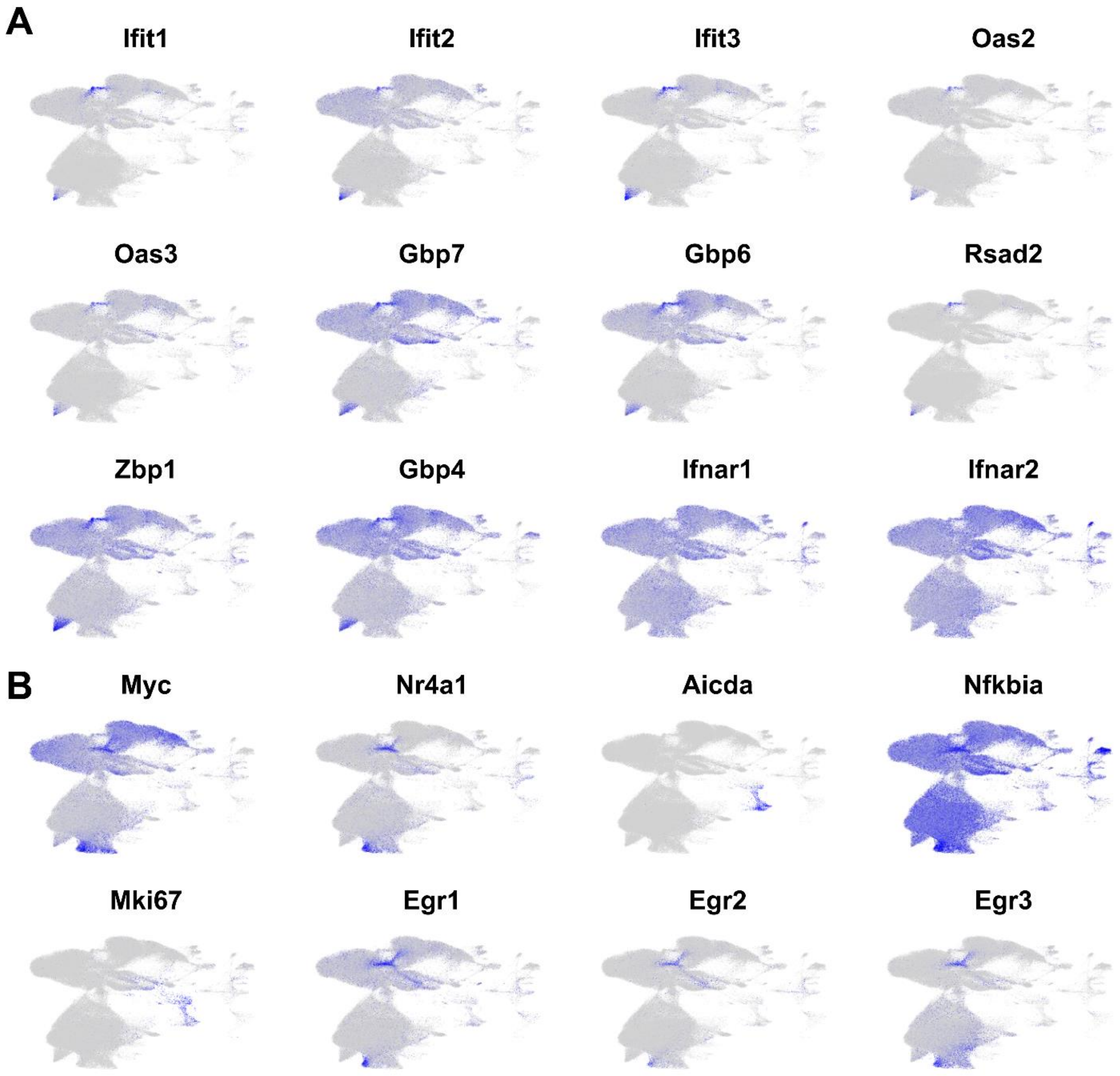


Fig S8: Gene expression heatmaps superimposed on UMAPs from dLN dataset. (A) Markers of antiviral response. Expression scale: <20$^{th}$ percentile = gray, >80$^{th}$ percentile = blue, 20-80 percentile → linearly scaled.

(B) Panel of immune signaling pathway markers. See text for details.

Description of supplementary tables:

Table S1: List of cell surface (A) and transcript (B) markers across cell clusters. Both positive and negative markers are catalogued. For the cut-offs used in determining the markers see Methods.

Table S2: Number of cells in each cell-cluster, across timepoints (D0, 1, 3, 5, 7, 30), exposure status (naïve, VIPEX), vaccination status (YF17D, InYF) and sex (M, F), from different tissues: (A) dLN, (B) spleen, (C) PBMC, (D) ndLN. Dint: internal control VIPEX.

Table S3: List of differentially expressed genes from clusters. Accompanies tissue, timepoint, vaccine, sex, gene name, cluster information, apart from log2FC and adjusted p-values.

Table S4: Gene groups across timepoint, sex and tissues. Number of genes in each group and member gene-cluster entries are listed.

Table S5: Fold change data with kinetic pattern annotations across timepoint, sex and vaccination status for all DEG-cluster entries from responding tissues. (A) dLN, (B) PBMC, (C) spleen. For pattern ids see figures S5 and S6.

Table S6: Enriched motifs in the promoters of genes from enriched cell-clusters within kinetic-patterns. The patterns discussed in the main text are prioritized. (A) dLN, (B) spleen, (C) spleen. The pattern and cluster enrichments are represented with ':'; pattern:cluster. The p-values and odds ratios of enrichments can be found in Figure S7A-C.
