## Supplementary Tables (Chatterjee et al) for "Kinetic patterns of single cell gene expression discriminate between the murine cellular responses to live attenuated and inactivated Yellow Fever vaccines": Table S6.pdf

**A**

| Pattern:cluster | Genes | Motif | Motif name | P-value | Adjusted p-value |
|---|---|---|---|---|---|
| LNI:τ6 | Igfbp4, Arid5a, Ndrg3, Skap2, Etv6, Jak3, Cdkn2d, Arid5b, Sbno2, Gngt2, Ly6e, Ly6a, Apobec3, Flot1, Zbp1, Crlf2, Abi3 | 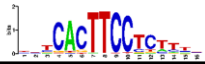 | Spib | 1.01887e-05 | 0.0076007702 |
| LNI:τ13 | Gbp5, Gbp2, Gbp4, Tgtp2, Aopep, Iigp1, Cd274, Cd74 | Same as above | Spib | 0.000116397 | 0.086832162 |
| LNI:d1 | Fcer1g, Hspa5, Hnrnpa3, Il1b, Cebpb, Ctsz, Prdx1, Ctsc, Pkm, Lilrb4a, Srgn, Lrp1, Plek, Kdm6b, Cd68, Eif4a1, Pfn1, Nme1, Atp5g1, Hif1a, Tgfbi, Ctsb, Atp6v0c, Myl12a, Junb, Eif5a | 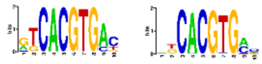 | Bhlhe41, Bhlhe40 | 1.28222e-05, 2.488e-05 | 0.0095653612, 0.01856048 |
| LNI:d2 | Arpc2, Actr3, S100a10, Wdr1, Ahcyl2, Fah, Ifitm2, Anxa2, Myl6, Pfn1, Gngt2, Myl12a, Cfl1, Flna, Hsp90aa1 | 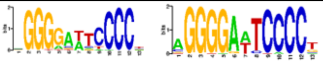 | Nfkb2, Nfkb1 | 3.23585e-05, 8.76699e-05 | 0.024139441, 0.0654017454 |
| LNII:b8 | Prkcb, Hdac9, Cd40, Emp3, Swap70, Grap, Lyst, Sh3bp5, Ms4a4c, Fcmr, Fam107b, Pde7a, Tmem131l, Rap1gds1, Bach2, Abca1, Akna, Marcksl1, Spen, Cnot6l, Rilpl2, Bmt2, Chd4, Fam169b, Cmip, Cyba, Reep3, Cdk17, Ikzf3, Rara, Nfkbia, Lpcat1, Cd180, Map3k1, Dleu2, Egr3, St13, Ccdc50, Btla, Ltb, Lbh, Rasgrp3, Eml4, Dmxl1, Bcor, Pikfyve, Tmem163, Ier5, Mndal, Akt3, Zbtb18, Itpkb, St8sia6, Vav2, Rapgef1, Itga4, Trim44, Dusp2, Zfp831, Il12a, Cd2, Nfkb1, Tspan5, Hivep3, Macf1, Id3, Atad3a, Cdk6, Sh3bp2, Tbc1d1, Tec, Sept11, Rasgef1b, Arhgap24, Kdm2b, Ncf1, Irf5, Iqsec1, Irak2, Ptpn6, Aebp2, Etnk1, Tgfb1, Spib, Arhgap17, Colgalt1, Piezo1, Maml2, Icam1, Fli1, Kmt2a, Myo1e, Shisa5, Hivep2, Marcks, Gadd45b, Osbpl8, Stk10, Dnmt3a, Ywhaq, Ifi27l2a, Traf3, Hivep1, Fam167a, Mycbp2, Pvt1, Kmt2d, Ttc3, Snx9, Ezr, Chd1, Ift140, Nfkbie, Foxp4, Tgif1, Egr1, Nfatc1, Nfkb2, Rp2, Tbl1x, Sms, Pard3b, Creb1, Dgkd, Bcl2, Btg2, 4930523C07Rik, Rcsd1, Fcrla, Tagln2, Ifi203, Tank, Cd44, Helz2, Dennd4b, Notch2, Cd72, Mier1, Adap1, Bri3, Bhlhe40, Zfp296, Rras2, Il21r, Nfat5, Gse1, Peak1, Dennd4a, Tgfbr2, Zbtb7a, Elk3, B4galnt1, Stat6, Plek, Sptbn1, Adam19, Kdm6b, Med11, Pecam1, Elmo1, Zswim6, Blk, St3gal1, Slc38a2, Plcl2, Gypc, Ehd1, Ahnak, Xiap, Ifi208, Fnbp1, Kynu, Tet3, Cdyl2, Irf2bp2, Tbx21, Ccr7, H2-K1, Aff3, Pgap1, C130026I21Rik, Ivns1abp, Rgl1, Ephx1, Vim, Otud1, Arhgap15, Cobll1, Cep250, Zbtb10, Tnik, Skil, Fcrl1, S100a10, Mcl1, Vav3, Lef1, Mllt3, Rell1, Oasl1, Orai1, Stx2, Rabgef1, Hip1, Slc7a1, Samd9l, Mat2a, Itpr1, Tmcc1, Fkbp4, Clec2i, Kras, Myadm, Pglyrp1, Relb, Apoe, Nfkbid, Gramd1a, Gm26827, Akap13, Ppfibp2, Gm16201, Irf7, Ier2, Gm42031, Casp4, Ubash3b, Cxcr5, Pml, Myo9a, Gm39383, Ipcef1, Stx11, Tnfaip3, Cd24a, Jmjd1c, Icosl, Mob3a, Atp2b1, Nab2, Gm16229, Gdf11, B3gnt2, Rel, Rapgef6, Ksr1, Kcnh4, Dusp3, Rnf213, Gpr65, Gpr132, Hist1h2bc, Cmah, Ssbp2, Mast4, Parp8, Arhgef3, Prkcd, Irf9, Kctd12, Klf10, Trps1, Zhx2, Myc, Ly6d, Ly6a, Tcf20, Litaf, Tfrc, Zdhhc14, E430024P14Rik, Tagap, Tnf, Lta, H2-Q6, H2-Q7, Prkce, Socs5, March3, Tcf4, Neat1, Sh3pxd2a, Cybb, Dock11, Gpr174, Sh3kbp1, Clasp1, Rabgap1l, Rc3h1, Il2ra, Traf1, Gabpb1, Hck, B4galt5, Maml3, Bcl10, St6galnac3, Zc3h12a, Phtf2, Card11, Trim24, Gimap7, Pde1c, Wnk1, Slc7a6, Plekho2, Inka1, Ifngr1, Map3k5, Srgn, Uhrf1bp1l, Tmcc3, 9930111J21Rik2, Med13, Ifrd1, Susd6, Nln, Slc25a37, Fndc3a, Mapk11, Nr4a1, Snn, 2010309G21Rik, Nfkbiz, Lst1, Wdr43, Fbxo11, Gramd3, Chka, Ddx3x, Hspd1, Nfkbib, Cmtm7 | 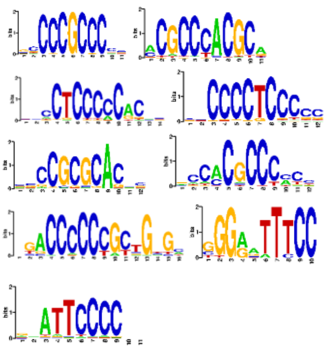 | Klf15, Egr2, Wt1, Znf148, Zbtb14, Sp9, Zic5, RelA, Rel | 9.76439e-21, 3.6826e-19, 8.5071e-18, 9.80227e-18, 1.90848e-16, 5.5448e-15, 1.23212e-14, 4.35088e-12, 1.15144e-06 | 7.28423494E-18, 2.7472196E-16, 6.3462966E-15, 7.31249342E-15, 1.42372608E-13, 4.1364208E-12, 9.1916152E-12, 3.24575648E-9, 0.00085897424 |
| LNIII:d1 | Il1rn, Sdc4, Ifitm2, F10, Msr1, Cstb, Ms4a6d, Gda | 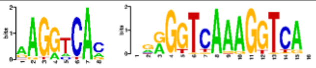 | Nr4a2, Ppard, | 5.98932e-05, 6.12604e-05 | 0.0446803272, 0.0457002584 |
| LNIV:τ12 | Stat1, Zbp1, Gbp2, Isg15, Gbp4, Irf7, Tgtp2, Ifi47, Irf1, Igtp, Irgm2, Fdft1, Iigp1, Irgm1, Serpina3g, Cxcr3 | 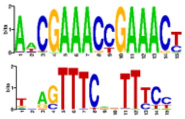 | Irf9, Stat1:Stat2 | 2.17058e-27, 2.70947e-12 | 1.61925268E-24, 2.02126462E-9 |
| LNIV:τ10 | Tgtp2, Ly6a, Stat1, Gbp7, Gbp2, Gbp4, 9330175E14Rik, Irgm1, Ifi47, Irf1, Igtp, Irgm2, Ccl5, Cd274, Gbp6 | Same as above | Irf9, Stat1:Stat2 | 2.65872e-16, 3.70516e-11 | 1.98340512E-13, 2.76404936E-8 |
| LNIV:b12 | Gbp2, Gbp4, Irgm1, Ifi47, Irf1, Socs1, Iigp1, Cd274, Plac8, Serpina3f, Tuba1b | Same as above | Irf9, Stat1:Stat2 | 2.32398e-11, 3.35517e-06 | 1.73368908E-8, 0.00250295682 |
| LNV:b10 | Pkib, Samsn1, Snx29, St7, Rflnb, Scn4a, Klhdc2, Gadd45g, Socs1, Pltp | 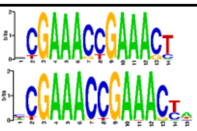 | Irf8, Irf4 | 2.88377e-06, 2.80522e-05 | 0.00215129242, 0.0209269412 |
| LNVII:τ4 | Itga4, Gm2682, Flna, AW112010, Jun, Klf2, Ier5, Tubb4b, Pag1, Txnip, 5031425E22Rik, Znrf2, Adgre5, Piezo1, Sorl1, Tgfbr2, Ppm1h, Arhgap5, Klf6, Zbtb20, H2-Q6, H2-Q7, Smad7, Neat1, Diaph2, Mitd1, Il18r1, Nabp1, Plekhm3, Tuba4a, Arl4c, Bcl2, Ptpn4, Cd55, Dstyk, Fasl, Tagln2, Ifi206, Mndal, Ifi203, Akt3, Capn2, Lamb3, St8sia6, Bmyc, Lypd6b, Tanc1, Rbms1, Ube2l6, Usp50, Slc20a1, Lbp, Slc17a9, Lmna, Atp8b2, Hist2h4, Slc25a24, S1pr1, Gbp5, Gbp7, Dnajb4, Ddx58, Reck, Cntln, Plin2, Acer2, Usp24, Kif1b, Tnfrsf25, Ski, Phtf2, Sept11, Abcg3, Gbp8, Gbp6, Tpst2, Oas1a, Vps37b, Zfp113, Slc7a1, Fry, Gm20559, Mdfic, Nod1, Herc6, Rabac1, Zbtb32, Nkg7, Gm45552, Hdgfl3, Trim30d, Gm20663, Gm44777, Xylt1, Rexo5, Hirip3, Ifitm10, Arhgef18, Rasa3, Tdrp, Ddx60, Bst2, Nlrc5, Sipa1l2, Ccdc15, Crtam, Tent5a, Nt5e, Mthfd1l, Esr1, Ust, Aig1, Gm26740, Scml4, Gm34006, P4ha1, Mcu, Cabcoco1, Ggt5, Hsp90b1, Atp2b1, BC048403, Itga7, Insyn2b, Pttg1, Irgm1, Gm5431, Gm12185, 9930111J21Rik1, Tgtp2, Jade2, Pik3r5, Rtn4rl1, Nufip2, Slfn5, Mrm1, Dhx40, Zfp652, Tbx21, Rara, Ubxn2a, Immp2l, Gtf2a1, Dglucy, Gpr68, Hsp90aa1, Ppp1r13b, Tasor2, Hist1h4c, Ripor2, Hivep1, Ssbp2, Arsb, Iqgap2, Naip2, Adamts6, Gm48321, Nr1d2, Ankrd28, Lats2, Phf11c, | 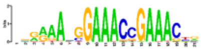 Only Irf3 motif shown, rest same as above | Klf15, Stat1:Stat2 Irf3, Znf148, Sp9 | 1.39409e-08, 5.54719e-06, 1.18043e-05, 1.62367e-05, 3.59205e-05 | 1.03999114E-5, 0.00413820374, 0.0088060078, 0.0121125782, 0.026796693 |

### A cont.

| | | | | | |
|---|---|---|---|---|---|
| LNVII:$\tau$16 | Sptssa, Mgat5, Coro2a, Tmem176a, Jaml, Atp2b1, Emb, Mycbp2, Il7r, Arfgef1, Il18r1, Il18rap, Stat1, Ikzf2, Fam124b, Dgkd, Ramp1, Bcl2, Ptpn4, Sox13, Camsap2, Rnasel, Rabgap1l, Atf6, Slamf1, Ifi203, Cep170, Sdccag8, Akt3, Smyd3, Smyd2, Il2ra, Nebl, Spopl, Psd4, Bmyc, Rapgef1, Fnbp1, Mbd5, Myo3b, Pde11a, Trp53i11, Trim44, Mga, Eid1, Smox, Gpcpd1, Tasp1, Sdc4, Eya2, Pde7a, Tnik, Phc3, Il2, Golim4, Lmna, Gm38411, Rorc, Lingo4, Tdrkh, Txnip, Hipk1, Ptpn22, Slc25a24, Prss12, Lmo4, Hs2st1, Tmem64, Ankrd6, Akirin2, Aqp3, Trim14, Zfp462, Cdk5rap2, Megf9, Dennd4c, Usp24, Tut4, Eps15, Cited4, Camk2n1, Iffo2, Agtrap, Kif1b, Tnfrsf25, Espn, Ski, Fgl2, Kmt2c, Crmp1, Fryl, Clock, Sept11, Antxr2, Rasgef1b, Gbp8, Lrrc8b, Rasa4, Glcci1, Mdfic, Lncpint, Tmem176b, Osbpl3, Jazf1, Il23r, Capg, Exoc6b, Mitf, Eif4e3, Il17re, Plxnd1, Wnk1, Atn1, Tnfrsf1a, Fkbp4, Klrk1, Smim10l1, Dusp16, Itpr2, Resf1, Tshz3, 1600014C10Rik, Ptov1, Kcnc1, Nav2, Igf1r, Slco3a1, Tm6sf1, Arnt2, Arrb1, Pgm2l1, Fchsd2, Arntl, Pde3b, Sh2b1, Itgal, Adam12, Cd163l1, 5830411N06Rik, Olfr60, Cers4, Irs2, Rab11fip1, Dlc1, D130040H23Rik, Lrrc25, Mast3, Chd9, Lpcat2, Nlrc5, Rnf166, Kcnk1, Pard3, Sesn3, Icam1, Cdon, Sorl1, Cbl, Zbtb16, Peak1, Adpgk, Coro2b, Smad3, Dennd4a, Rora, Nedd4, Lysmd2, Gclc, Sh3bgrl2, Rab6b, Acpp, Gpx1, Map4, Clasp2, Tgfbr2, Cxcr6, Phactr2, Ifngr1, Raet1e, Gm26740, Cdk19, Sesn1, Spock2, Psap, Chst11, Cry1, Hsp90b1, Socs2, Nudt4, Ppp1r12a, Kcnmb4, Irak3, Ppm1h, Ptges3, Znrf3, Rtn4, Dock2, Insyn2b, Adam19, Tcf7, Fstl4, Sept8, Rad50, Ncor1, Chd3, Cpd, Abhd15, Slfn8, Rnf43, Msi2, Scpep1, Mmd, Hlf, Fam117a, Zfp652, Ikzf3, Thra, Nr1d1, Tex2, Pitpnc1, Rnf213, Sdc1, Odc1, Ywhaq, Hectd1, Arhgap5, Syne2, Sipa1l1, Gpr65, Serpina3g, Ckb, Zmynd11, Actn2, Edaradd, Lyst, Serpinb1a, Serpinb6a, Pxdc1, Mcur1, Card19, Slc12a7, Cast, Ssbp2, Parp8, Nr1d2, Dlg5, Chdh, Rnase4, Ltb4r1, Wdfy2, Blk, Xkr6, Kif13b, Fndc3a, Rgcc, Mbnl2, Selenop, Cdh10, Retreg1, Ank, Trio, Ubr5, Sntb1, Fam83a, Zhx1, Tmem65, Grap2, Bik, Cerk, Rarg, Pcbp2, Crebbp, Dexi, Socs1, Comt, Arhgap31, Zbtb20, Abi3bp, St3gal6, App, Tiam1, Ttc3, Hmgn1, Synj2, Qk, Igf2r, Airn, Mmp25, Baiap3, Zfp871, Myo1f, Ly6g5b, H2-Q4, H2-Q6, H2-Q7, Gabbr1, Trerf1, Tbc1d5, Tnfsf14, Pja2, Crim1, Prkce, Ttc7, Mib1, Ttc39c, Galnt1, Ankhd1, Diaph1, Nr3c1, Prelid2, Ap3s1, March3, Slc27a6, Ldlrad4, Zfp516, Vegfb, Anxa1, Klf9, Pip5k1b, 9930021J03Rik, Uhrf2, Pten, Pi4k2a, Slc25a28, Dnmbp, Atrnl1, Fgf13, Pola1, Ar, Ftx, Gprasp1, Huwe1, AC149090.1, Arhgap21, Dnajc10, P2ry1, Ankrd13c, Ddx58, Vps13d, 5830444B04Rik, Gbp4, Dipk1a, Gm20559, Braf, Fam160a2, Xylt1, Zdhhc2, Ddx60, Bst2, Aagab, Plekho2, Myo1e, Appl2, Camk2b, Nudcd3, Irgm1, Olfr56, Tanc2, Lgals3bp, Immp2l, Lats2, Atad2, Tnrc6b, Cd96, Mex3c, Pacs1, Ctsw, Gab3 | 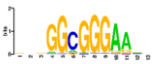 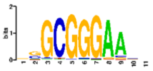 Motifs for E2f6 and Tfdp1 respectively. Rest are same as above. | Klf15, Zbtb14, Sp9, E2f6, Tfdp1 | 2.42977e-16,<br>7.61248e-14,<br>2.83907e-12,<br>1.61191e-11,<br>4.0544e-11 | 1.81260842E-13,<br>5.67891008E-11,<br>2.11794622E-9,<br>1.20248486E-8<br>3.0245824E-8 |

**B**

| Pattern: cluster | Genes | Motif | Motif name | p-value | Adjusted p-value |
|---|---|---|---|---|---|
| SPI:ν1 | Ptpn18, Ankrd44, Cxcr4, Prdx6, Creg1, Atf3, Fam107b, Atp5c1, Gsn, Zc3h15, Cst3, Maml3, Golim4, Cd101, Sec61b, Elp1, Ptp4a2, Srpk2, Ost4, Dynll1, Taok3, Ncf1, Mdh2, N4bp2l1, Cycs, Rab43, Clec4a2, Clec12a, Plbd1, Lamp1, Myo9b, Ndufb7, Tecr, Pkn1, Cox4i1, Irf2bp2, Slc9a9, Utrn, Rwdd1, Psap, Palm, Tle5, Plxnc1, Atp2b1, Lyz2, Cd63, 4930438A08Rik, Rap1gap2, Limd2, Cd300a, Cd300c2, Rhob, Arhgap5, Nin, Calm1, Dglucy, Adssl1, Gpr141, Gm34084, Ninj1, Mctp1, 5430425K12Rik, Gm36161, Sap18, Itm2b, Mtdh, Ndufa6, Krt80, Klhl6, Mpc1, Myo1f, Lst1, Ankrd12, Myl12a, Cebpz, Sec11c, Mbp, Rnaseh2c, Fermt3, Polr2g, Gnaq, Pgam1, Tmsb4x, Arfgef1, Neurl3, Inpp5d, Dgkd, Lpgat1, Apbb1ip, Itga4, Csnk2a1, Bri3, Polr1d, St3gal5, Grcc10, Tgfb1, Tm6sf1, Acsl1, Man2b1, Plcg2, Hexa, Gm2a, Atox1, Vps13b, Sh3bp1, Alcam, Ezr, Unc93b1, Fau, Slc25a5, Adgre5, Pla2g7, Mpeg1, Kpna4, Tut4, Ppt1, Ptafr, Tmem140, Xylt1, Rasa3, Aprt, Cox5a, Glipr1, Rassf3, Dock2, Slc43a2, Rsad2, Dleu2, Retreg1 | 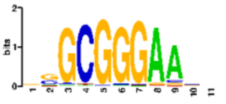 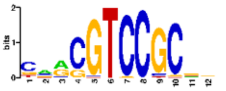 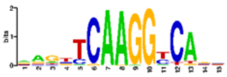 Motifs for Klf6, Hinfp and Nr5a2 are shown. Rest are same as in S13 P. | Klf15,<br>Klf6,<br>Sp9,<br>Hinfp,<br>Nr5a2 | 1.26196e-07<br>2.92131e-06<br>1.39545e-05<br>4.05865e-05<br>9.1781e-05 | 9.4142216E-5<br>0.002179297<br>0.010410057<br>0.030277529<br>0.068468626 |
| SPII:μ1 | Ptma, Fcer1g, Hspa5, Hnrnpa3, Rpn2, Rpn1, Ifitm2, St3gal4, Lilrb4a, Srgn, Sarnp, Eif5a, Lgmn, Lcp1, Smdt1, Elob, Trem3, Fkbp2, Ap1s2, Prdx1, Clec4a3, Gm15987, Arhgdib, Ldha, Ppia, Grina, Ap2m1, Gngt2, Msn, Tmem258, S100a11, Cfl1 | 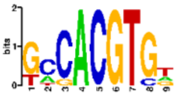 | Creb3l2 | 6.59558e-05 | 0.0492030268 |
| SPIII:ν1 | Arid5a, Rnasel, Qsox1, Smox, Ppp1r3d, Fabp5, Slc2a1, Steap4, Fbxl5, Ube2h, Tarm1, Mrgpra2b, Il4ra, Ifitm2, Ifitm1, Atp11a, Ppp1r3b, Klhdc4, Casp4, Entpd3, Lilr4b, Lilrb4a, Smpdl3a, Osm, Adam19, Atp6v0a1, Socs3, Hif1a, Rdh12, Zfp36l1, Spata13, Azin1, Mirt2, Csf2rb, Apobec3, Tspo, Atxn10, Lpp, Il1rap, App, Ier3, Lrg1, Mta3, Mbd2, Il13ra1, Gk, Bcl2l1, Tlr2, Mgam, Sipa1l1, Cd14 | 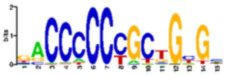 | Zic3 | 7.27246e-05 | 0.0542525516 |
| SPIV:τ5 | Ucp2, Ahnak, Klrd1, Adgre5, Gm21887, Itga4, Rbm3, Sp100, Samhd1, S100a6, S100a10, Laptm5, Cd52, Selplg, Actb, Zyx, Cd8b1, Gapdh, Gmfg, Gm45552, Gm4070, Coro1a, Pycard, Ctsd, Lsp1, Hmgb2, Thy1, Ccr5, Itgb2, Eef2, S1pr4, Dctn2, Anxa6, Grap, Ccl5, Armc7, Actg1, Ndufb1-ps, Epsti1, Lgals1, Rnaset2b, Gstp3, Tbc1d10c, Cfl1, Ms4a4b, Ms4a6b, Tmem163, Slamf7, Cd48, Vim, Ctsa, S100a13, Hip1, Arpc4, Arhgdib, Spn, Slc25a4, Itgb1, Abracl, Rap1b, Plek, Crip1, Serpinb6b, Glrx, Sh3bgrl, Fry | 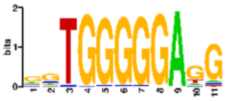 | Znf281 | 5.7823e-05 | 0.043135958 |
| SPV:τ14 | Dennd1a, Magi3, Tapt1, Cux1, Rmnd5a, Ptms, Lockd, Tcf12, Lgals9, Rcbtb2, Xrcc6, Desi1, Ano6, Slc38a2, Runx1, Fkbp5, Cyb5a, Agfg2, Itpr2, Mef2a, Mns1, Slc9a9, Xrn1, Ifngr1, Myb, Stat5b, Rgcc | 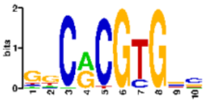 | Hes1 | 8.76148e-05 | 0.06536064 |
| SPVI:τ14 | Glcci1, Ppm1h, Atp10a, Znrf1, Endou, Ldlrad4 | 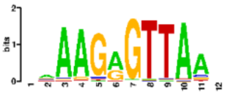 | Znf652 | 5.36226e-05 | 0.0400024596 |

### B cont.

| Pattern: cluster | Genes | Motif | Motif name | p-value | Adjusted p-value |
|---|---|---|---|---|---|
| SPVII:τ14 | Btg2, Lef1, Aff1, Clk3, Zfp280d, Osbpl8, H2afv, Rb1, Ly6e, Chd1, Zeb1, Ppp1r14b, Aspm, Rcsd1, Pip4k2a, Ckap5, Emc7, Ect2, Tmem50a, Hp1bp3, Nsd2, Chchd3, Ezh2, Tax1bp1, Edem1, Ncapd2, Atf7ip, Klf13, Prc1, Nsd3, Neil3, Ctcf, Cdkn2d, Kif23, Ccnb2, Fbxo5, Bsg, Top2a, Mxd3, Dlgap5, Dnajc3, Rad21, Zfp148, Dock11, Thoc2, Sh3kbp1 | 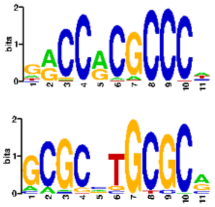 Motifs for Klf3 and Nrf1 are shown. | Klf3<br>Nrf1<br>Hinfp<br>Sp3 | 3.50879e-06<br>5.52553e-06<br>1.02338e-05<br>3.22694e-05 | 0.0026175573<br>0.004122045<br>0.0076344148<br>0.0240729724 |

## C

| Pattern: cluster | Genes | Motif | Motif name | p-value | Adjusted p-value |
|---|---|---|---|---|---|
| PBI:ν1 | Ptpn18, Plekhb2, Neurl3, Ankrd44, Plekhm3, Cab39, Inpp5d, Arl4c, Cxcr4, Kif21b, Ptgs2os2, Xpr1, Mrps14, F5, Creg1, Pfdn2, Iars2, Atf3, Lpgat1, Fam107b, Celf2, Uap1l1, Tbc1d13, Sh2d3c, Mvb12b, Gsn, Arhgap15, Itga4, Chac1, Slc20a1, Vps16, Cst3, Samhd1, Tti1, Ncoa3, Ctsz, Ppdpf, Sirpb1a, Mbnl1, Kpna4, Chtop, Cd101, Csf1, Sars, Snhg8, Manba, Sec61b, Elp1, Gng10, Macf1, Oscp1, Marcksl1, Ptp4a2, Laptm5, Man1c1, Syf2, Srsf10, Pithd1, Srpk2, Prkag2, Smim14, Stap1, Abcg3, Mlec, Dynll1, Tpcn1, Kmt5a, Bri3bp, Chchd2, Gusb, Orai2, Gm36551, Pilrb2, Daglb, Bri3, N4bp2l1, Exoc4, Nod1, St3gal5, Vamp5, Aup1, Pcbp1, Irak2, Tatdn2, Hnrnpf, Slc6a13, Ptpn6, Atn1, Scnn1a, Clec12a, Plbd1, Fosb, Atp1a3, Tgfb1, Bax, Nucb1, Emp3, Svip, Uvrag, Arrb1, Igsf6, Chst15, Lsp1, Lamp1, Rasa3, Mcph1, Tex15, Acsl1, Zfp868, Uba52, Isyna1, Haus8, Tecr, Pkn1, Nfix, Lyl1, Man2b1, Plcg2, Hsdl1, Rnf166, Tmem123, Slc37a2, Zpr1, Acat1, Hexa, Anp32a, Dennd4a, Bcl2a1b, Morf4l1, Pik3cb, Apeh, Ngp, Gpd1l, Higd1a, Pcmt1, Utrn, Psap, Ube2g2, Palm, Nfic, Hsp90b1, Plxnc1, Dusp6, Nap1l1, Slc16a7, Ptges3, Sptbn1, Rmnd5b, Jade2, Hint1, Gm2a, Atox1, Galnt10, Nlrp3, Pigl, Ctc1, Per1, Eif5a, Nup88, Rap1gap2, Slc43a2, Abhd15, Rhot1, Gngt2, Limd2, Cd300a, Cd300lb, Cd300c, Cd300c2, Srsf2, P4hb, Anapc11, Rhob, Rock2, Arhgap5, Mia2, Nin, Dhrs7, Acot1, Dglucy, Chrm3, Gpr141, Elmo1, Hist1h1c, Gfod1, Gm34084, Ninj1, Zfp346, Mctp1, Emb, Vdac2, Lrmda, Nisch, Timm23, Glud1, Tmem260, Sap18, Dock5, Itm2b, Gtf2f2, Dgkh, Selenop, Mtdh, Vps13b, Ankrd46, Ndrg1, Arhgap39, Sh3bp1, Ndufa6, Parvg, Dip2b, Nr4a1, Csad, Rarg, Coro7, Alg1, Klhl6, Alcam, Gbe1, Sod1, Scaf4, Rnaset2b, Mpc1, Tmem8, Myo1f, H2-K1, H2-Ke6, Lst1, H2-D1, Gm7030, Tnfrsf21, Pla2g7, Slc35b2, Vegfa, Adgre1, Ankrd12, Cdc42ep3, Gm19696, Calm2, Mapre2, Etf1, Txnl1, Sec11c, Pmaip1, Impa2, Seh1l, Smad2, Pstpip2, Mbp, Ndufs8, Unc93b1, Gstp1, Ccs, Drap1, Rnaseh2c, Map4k2, Fermt3, Cpsf7, Gnaq, Frat2, Eif3a, Grk5, Slc25a5, Cox7b, Txlng, Hnrnpu, Maml3, Txnip, Fgl2, Fxyd5, AI467606, Itgal, Adgre5, Jaml, Peak1, Slc9a9, Bcr, Hist1h4i, Hist1h2bc, Mpeg1 | 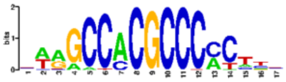 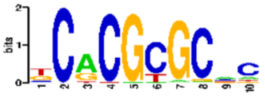 Motifs for Sp4 and Tcfl5 are shown. Rest of the motifs are shown in previous figures. | Klf15, Sp4, Sp3, Tcfl5, Zbtb14, Hes1, Nrf1, Zic5 | 7.83041e-13, 1.18387e-09, 1.91162e-09, 2.36375e-09, 2.16037e-08, 1.30238e-07, 1.81439e-07, 2.21999e-06 | 5.84148586E-10, 8.8316702E-7, 1.42606852E-6, 1.7633575E-6, 1.61163602E-5, 9.7157548E-5, 0.000135353494, 0.00165611254 |
| PBIII:ν1 | Arid5a, Rufy4, Hdac4, Pam, Rnasel, B4galt3, Lbr, Cers6, Morrbid, Smox, Sycp2, Ppp1r3d, Fabp5, Gyg, Tlr2, S100a6, Rhoc, Mov10, Gclm, Glipr2, Atp6v1g1, Rnf11, Slc2a1, Steap4, Fbxl5, Rbm47, Atp8a1, Arhgap24, Cux1, Gpr146, Alox5ap, Ube2h, Gadd45a, Klra2, Dusp16, Tarm1, Cd33, Asb7, Il4ra, Ifitm2, Mcemp1, Atp11a, Ppp1r3b, Mtus1, Crispld2, Klhdc4, Casp4, Cdkn2d, Ddx6, Dmxl2, Pde7b, Lilr4b, Lilrb4a, Smpdl3a, Srgn, Ggt5, Sbno2, Xbp1, Adam19, Wfdc21, Atp6v0a1, Socs3, Hif1a, Rdh12, Zfp36l1, Sipa1l1, Rab24, Spata13, Acod1, Azin1, Dgat1, Csf2rb, Apobec3, Prr13, Lpp, Il1rap, App, Ier3, Flot1, Lrg1, Mbd2, Gda, Atrnl1, Il13ra1, Gk, R3hdm4, Glrx, Lgals3, Atxn10, Chil1, Vim | 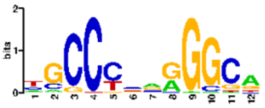 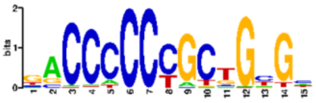 | Tfap2c, Zic3 | 2.00883e-06 5.4349e-06 | 0.00149858718 0.0040544354 |

#### C cont.

| Pattern:cluster | Genes | Motif | Motif name | p-value | Adjusted p-value |
|---|---|---|---|---|---|
| PBIII:μ1 | Fabp5, Ssr2, Krtcap2, Hmgn2, Fbxl5, Arhgap24, Tmem176b, Tmem176a, Clec4e, Ube2s, Bcl3, Ldha, Il4ra, Rnh1, Hp, Mmp8, Snx1, Manf, Myd88, Sbno2, Lta4h, Xbp1, Flot2, Socs3, Gcnt2, Nfil3, Prelid1, Tgfbi, Vcan, Mrpl52, Vps28, Csf2rb, Lgals1, Tspo, Sdf2l1, Flot1, Pla2g7, Mydgf, Ap3s1, Ms4a4a, Ms4a6d, Gda, Cfp | 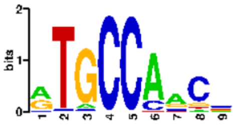 | Hic1 | 6.88065e-05 | 0.051329649 |
| PBIII:d1 | Il1b, Tpd52, Sirpb1c, Clec4a1, Clec4a3, Clec4a2, Siglece, Ctsc, Il4ra, Nupr1, Ifitm1, Ppp2cb, Gsr, Aprt, Myd88, Lilrb4a, Lgmn, Tgfbi, Ctsb, Cebpd, Msrb1, Aif1, Cdk2ap2, Fcgr2b, Serinc3, Abi3 | 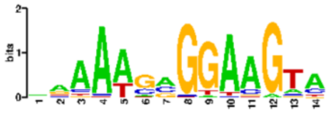 | Spic | 9.18363e-06 | 0.00685098798 |
